# A genomic dating tool resolves the origins of ancient Eurasian genomes

**DOI:** 10.1101/828962

**Authors:** U. Esposito, G. Holland, G. Alshehab, A. M. Dobre, M. Pirooznia, C. S. Brimacombe, E. Elhaik

## Abstract

Radiocarbon dating is the gold-standard in archaeology to estimate the age of skeletons, a key to studying their origins. Nearly half of all published ancient human genomes lack reliable and direct dates, which results in obscure and contradictory reports. Here, we developed the Temporal Population Structure (TPS), the first DNA-based dating method for ancient genomes ranging from the Upper Palaeolithic to modern-day samples and applied it to 1559 ancient Europeans, Asians, and Caucasus individuals and to 2117 modern worldwide individuals. We show that TPS predictions for dated skeletons align with their known dates and correctly account for kin relationships. The TPS-dating of poorly dated Eurasian samples resolves conflicts and sheds new light on disputed findings, as illustrated by four test cases. We discuss the phenotypic traits of the Time Informative Markers (TIMs) that underlie TPS.

**Summary:** TPS is a novel method to date humans from the Upper Palaeolithic to modern time from their DNA sequences.

Accurate dating is essential to the interepretation of paleogemonic data.. The gold-standard method in archaeology is radiocarbon dating^*1*^. However, a major limitation of radiocarbon dating is the high amount of collagen extraction (500 mg) involved in the process^*2*^. Consequently, half of all published ancient human genomes lack reliable and direct dates, which results in obscure and contradictory reports. Here, we present the Temporal Population Structure (TPS), the first genomic dating method for ancient genomes ranging from the Upper Palaeolithic to modern-day samples. We show that TPS predictions for 961 radiocarbon-dated Eurasian skeletons align with their known dates. We replicate these findings on 598 other Europeans, Asians and Caucasus individuals. Using kin-pairs, we demonstrate that TPS has produced more accurate results than radiocarbon and other dating. We show how our findings resolve conflicts and sheds new light on disputed findings as illustrated by four test cases. Finally, we discuss the phenotypic traits of the Time Informative Markers (TIMs) that underlie TPS. TPS is a novel dating technique, which can be used when radiocarbon dating is unfeasible or uncertain or to develop alternative hypotheses. TPS cannot be used for older (<14,000 years ago) samples, and its accuracy depends on the temporal and geographical breadth of radiocarbon-dated samples in the training dataset, though this limitation can be improved over time. Overall, TPS can improve the accuracy of archeological and paleogenomic studies.

## Background

Ancient DNA (aDNA) has transformed the study of human demographic history, allowing us to directly analyze patterns of past genetic variation rather than infer them *post factum* ^*3*^. The last few years have witnessed a conspicuous increase in the volumes of ancient skeletal DNA and studies attempting to trace their origins ^4^. Dating ancient remains is of key importance to the production of meaningful and reliable historical reconstructions, particularly in light of the growing medicalization of the field.

In the second half of the 20^th^ century, radiocarbon dating dramatically changed the field of archaeology ^5^ and became the gold standard to date ancient organic materials ^1^. Radiocarbon dating is based on the observation that living beings exchange ^14^C with their biosphere while alive and cease to do so when dead. At that point, their ^14^C atoms decay into ^14^N with a half-life of ∼5,700 y, whereas their ^12^C concentration remains constant ^6,7^. Thereby, assuming that the initial ratio of carbon isotopes in the biosphere remained constant over time, measuring the ^14^C to ^12^C ratio allows inferring the sample’s age. Over the years, improvements to the original method were made ^7^, including pre-treatment of the samples’ bones to eliminate contamination by recent carbon ^8^ and the introduction of accelerator mass spectrometry, which advanced the measurement of the decaying process ^9^. Improvements in the knowledge of Earth’s past environment and the quantification of reservoir effects and paleodiets have led to more accurate calibration curves of the past biosphere isotopes levels ^7,10-12^. For instance, the bones recovered from Repton (England) were associated with the Viking Great Army from 873-874 CE based on the archaeological context. However, early radiocarbon results predated some of them to the seventh and eighth centuries CE ^13^. Only a later radiocarbon analysis that considered the marine reservoir effects found that all dated remains are consistent with a single late ninth-century event, in line with the numismatic evidence ^14^. Overall, radiocarbon dating has been continuously improving in accuracy and reliability over the past decades since its proposal some 65 years ago.

A major limitation of radiocarbon dating is the high amount of collagen extraction (500 mg) involved in the process ^2^. To date, of the ∼2,600 ancient skeletons whose aDNA was successfully sequenced and published, less than 60% were radiocarbon-dated. The remaining skeletons have either been dated according to the archaeological materials found alongside the sample or remain undated. The subjective interpretation of skeletal data already led to misunderstandings on numerous occasions. For instance, a bone from the Darra-i-Kur cave in Afghanistan, initially assumed to be from the Palaeolithic (30,000 years BP [yBP]) ^15^ and often cited as one of the very few Pleistocene human fossils from Central Asia, was recently radiocarbon-dated to the Neolithic (4,500 yBP) ^16^. Similarly, one of the Brandysek site individuals (RISE569) was originally attributed to the Bell Beaker period (4,800-3,800 yBP) ^17^, but a later radiocarbon dating showed that the skeleton largely post-dated this culture (1,400-1,100 yBP) ^18^. Indeed, reevaluations of ^14^C calibration curves are not rare ^19^, not only different tissues produce different results, but labs may produce radiocarbon ages that differ up to and over 1,000 years ^2^. As misattributions can lead to erroneous conclusions, the uncertainty in the age of nearly half of the aDNA samples poses considerable risk of misinterpretation to the field, which calls into question their cost-effectiveness and overall usefulness.

Here, we present the Temporal Population Structure (TPS) tool, the first DNA-based dating method suitable for samples younger than 14,000 yBP. Compared to radiocarbon dating, DNA analyses require less material (x1/5) ^*20*^, which makes DNA-based dating methods more viable when there are limited materials. Thus, TPS can be used to directly date skeletal aDNA for which no radiocarbon date is available or as an independent validation approach for existing results. The only other DNA-based method for dating was developed for genomes older than 30,000-20,000 yBP and estimated the age of a sample from their Neanderthal ancestry ^*21,22*^. This technique, tested and demonstrated on five of the most ancient human genomes (45,000-12,000 yBP), becomes unstable for more recent dates ^*21*^ and is unsuitable for dating the vast majority of the ancient genomes younger than 10,000 yBP or non-Europeans genomes in general. By contrast, TPS was calibrated using existing Eurasians (14,000 yBP-present), which constrains its accuracy to the range of reliable radiocarbon dating.

As most allele frequencies in a population vary due to random mutations associated with demographic processes that change over time ^23^, we hypothesized that there exists a group of markers that exhibit substantially different allele frequencies between different time periods and can be used to estimate temporal trends. We termed these markers *Time Informative Markers* (TIMs). Conceptually, TIMs are reminiscent of ancient Ancestry Informative Markers (aAIMs) ^*24*^, except that they operate along the time axis. Random time-dependent events create unique allele frequencies in the TIMs that are characteristic of the historical period in which individuals lived, which we termed *temporal components*. Briefly, to date a sample, we first identified the temporal components from a database of ancient Eurasian genomes as the expression of their allele frequency variation over time (not over space). We then calibrated the temporal profiles using radiocarbon-dated ancient genomes and constructed a reference panel of temporal populations. TPS calculates the temporal components of the undated sample and compares them with those of the temporal components of the reference populations. TPS then dates the sample based on the radiocarbon dates of the temporal populations with the most similar dating component to those of the test sample.

### The Temporal Population Structure (TPS) model

We curated a database *DB*_*Anc*_ of 961 ancient Eurasian genomes (Fig. 1; Table S1) from the Late Upper Palaeolithic to the Anglo-Saxon Age (14,000-1,000 yBP). Of these, 602 (63%) were radiocarbon-dated (*DB*_*RD*_*) and* 321 (33%) were dated indirectly (usually from the archaeological context) (*DB*_*OD*_ ). The dates of the remaining 38 (4%) individuals were either unknown or unclear (*DB*_*UD*_).

**Fig. 1.**
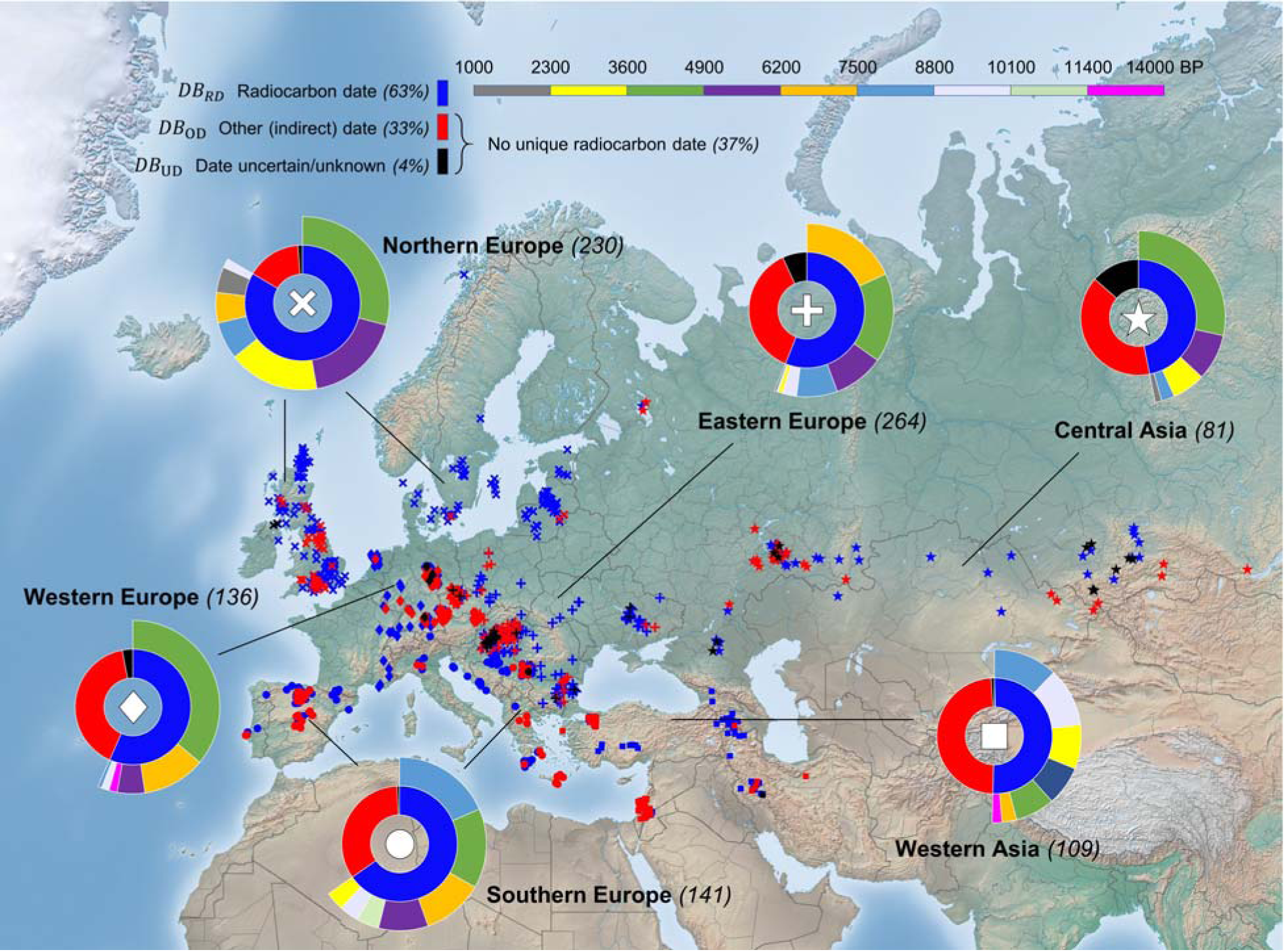
Location and dating of the 961 ancient individuals (D_Anc_) used in the initial calibration and testing of this paper. Symbols mark geographical macro-areas where samples were found. Colours represent their dating method (D_RD_ (blue), D_OD_ (red), and D_UD_ (black)). The inner circle of each sunburst shows the proportion of samples dated with different dating methods and the outer circle marks the distribution of radiocarbon dates for the local samples. Radiocarbon dating is divided into nine temporal bins of 1,300 years (top bar) covering the timeline of our database, excepting the oldest bin that is 2,600 years wide.

To construct the temporal components, we developed a cohort of ancient and modern genomes. For that, we merged a random subset of 300 *DB*_*Anc*_ genomes (Table S2) with a database *DB*_*Mod*_ of 250 individuals from Europe, Asia and Africa and applied *unsupervised* ADMIXTURE ^*25*^ with a various number of *K* components (Fig. S1). Five ancient and three modern temporal components captured the temporal trends of the combined database (Fig. S2). The allele frequencies of the temporal components were used to simulate the DNA of putative “populations” that represented typical genomes from various periods (Figs. S3-4; Table S2; Data S2).

A *supervised* ADMIXTURE analysis for *DB*_*RD*_ individuals against the temporal components (Fig. 2; Table S2) shows that each temporal component predominates a delimited time interval.

**Fig. 2.**
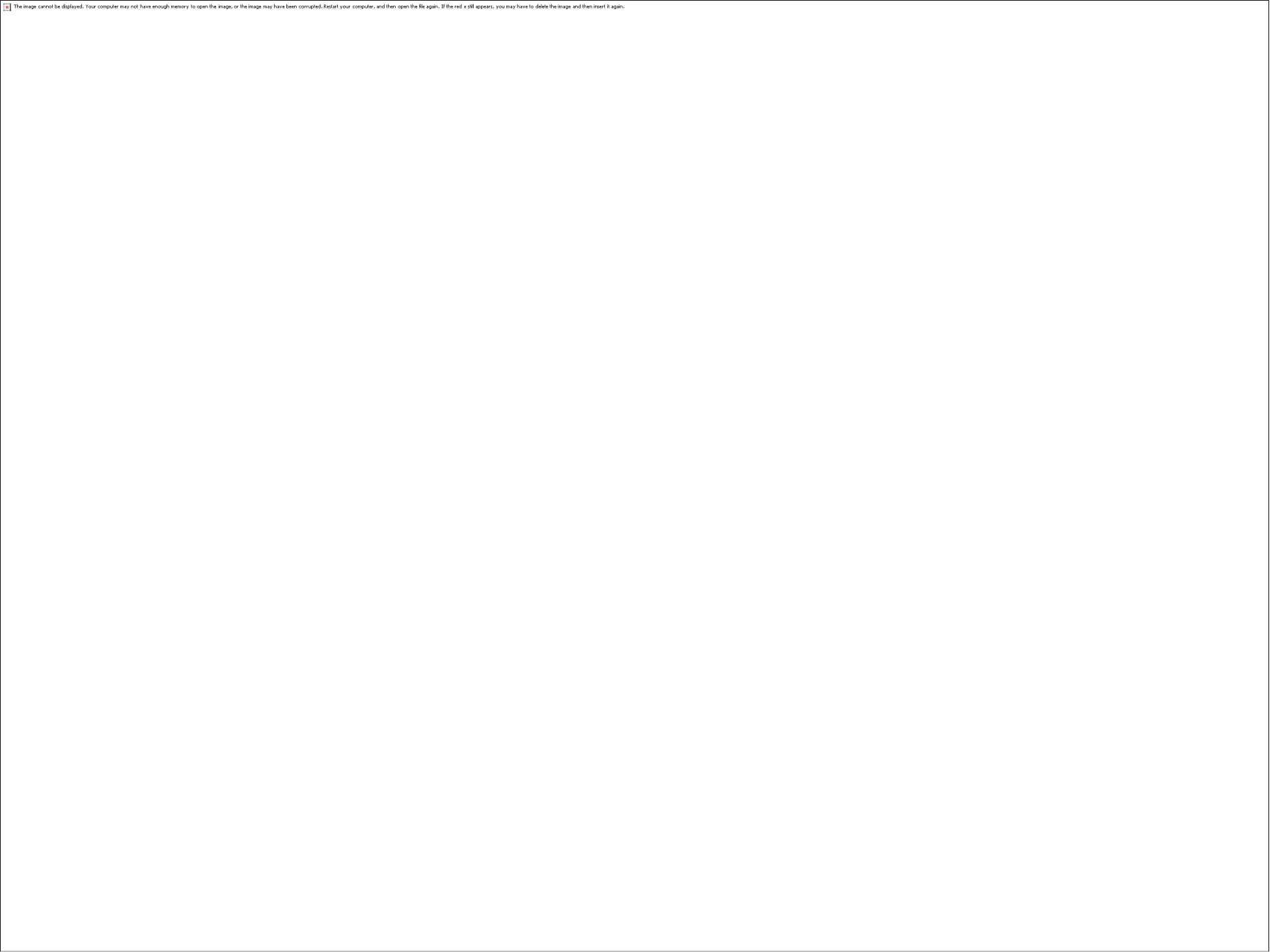
Ancient temporal components for the 602 radiocarbon-dated individuals (DB_RD_). Each vertical stacked bar represents an individual. Colours correspond to the five ancient temporal components. The three modern temporal components are meager for these samples and were omitted for coherency. Individuals are sorted by their age (note that the time x-axis is not linear). All the components are reported in Table S2.

The TPS reference panel consists of radiocarbon-dated individuals whose date was predicted well by TPS in a randomized setting. To identify those individuals, we assigned *DB*_*RD*_ individuals to non-overlapping bins of 500 years. TPS was applied 500 times to each individual. In each run, a random set of 400 individuals representing the diversity of all temporal profiles was selected and used to construct a reference panel of temporal populations. TPS then calculated the Euclidean distances between the temporal components of the test individual, which was not included in the reference panel, and those of the reference individuals and identified the three closest matching temporal individuals. The final date is obtained from the linear combination of the dates of these three individuals with weights proportional to the inverse of their Euclidean distances (Supplementary Materials). Based on these data, TPS then reports the average date and error estimate. By the end of this procedure, all predicted TPS dates per each individual were averaged across all runs and individuals whose averaged TPS prediction deviated from their radiocarbon date by over 1,000 years were discarded from the reference panel. The final TPS reference panel comprised of 496 ancient individuals. Reference individuals with comparable dates but different geographical locations exhibited a high similarity than samples from different time periods, regardless of their burial location (Fig S5).

### Identifying Time Informative Markers (TIMs)

To identify the genomic markers that underlie the ancient temporal components employed by TPS, the components were sorted from oldest to youngest (Fig. S6), creating a temporal variation profile of allele frequencies for every SNP. A time-series analysis (Supplementary Materials) identified 62,371 out of 150,278 SNPs whose frequency either decreased or increased over at least 3,000 years (Fig. S7), which we termed Time Informative Markers (TIMs) (Table S3) and 24,311 non-TIMs whose allele frequency exhibited little or no variation over time. Unsurprisingly, most of the TIMs (53%) are intronic and 13% are intergenic ^*26*^; 52% of the coding variants are missense (Fig. S8). Due to the high missingness of the dataset, to avoid omitting samples (Supplementary Materials), we used the entire SNP set for the remaining analyses.

### Evaluating the accuracy of TPS

Next, we compared the radiocarbon dates for the individuals in the reference panel with their TPS predictions obtained through the leave-one-out individual testing procedure (Fig. 3). The two dating measures were highly correlated (two-sided *t-test, N*=496: *r*=0.95, *95% CI*: [0.947, 0.963], *p*=1.4e-263) and in a close alignment to the ideal bisecting line *y = x* (Fig. 3A). To gauge the reliability of TPS predictions, we defined the accuracy per individual as the absolute difference between the TPS result and the radiocarbon date. The median accuracy for this testing procedure was 324 years and 68% of the individuals were assigned a TPS date within 500 years from their radiocarbon date. Similarly, the 95% error intervals of TPS (Supplementary Materials) and the confidence interval radiocarbon dates overlapped for 69% of the 496 individuals (Table S4). Only 4% of the individuals were TPS-dated over 1,000 years from their radiocarbon date (Fig. 3B; Table S4). The general uniformity in the distribution of the accuracy across the different time periods suggests the absence of temporal biases towards any particular period, with the exception of the oldest samples for which performance is below average, most likely due to the small number of samples. Overall, TPS results matched the radiocarbon dates (Fig. 3C), with exceptionally high accuracy in the 168 Northern Europeans (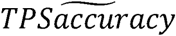=269 years). We found no statistical difference in the mean distribution of TPS’s accuracy between the major consecutive periods (two-sided *t-test, N*_*Mesolithic*_=63, *N*_*Neolithic*_=208, *p*=0.065; *N*_*Neolithic*_=208, *N*_*Copper*_=136, *p*=0.66; *N*_*Copper*_=136, *N*_*Bronze*_=121, *p*=0.19). TPS predictions of same-country individuals also did not cluster around a single time period and were spread over the timeline following their radiocarbon dates, confirming that the temporal components represent temporal rather than geographical variation (Fig. S9). TPS accuracy was not correlated with the genomic coverage (Fig. S10), suggesting that TPS is robust to missing SNPs (within the set limit of including samples with over 15k SNPs). TPS results for the non-TIMs were comparable to the results of two null models (Figs. S11-2). By contrast, TPS results for the TIMs (Fig. S13) yielded similar results to those of the full set of SNPs (Fig. 3).

**Fig. 3.**
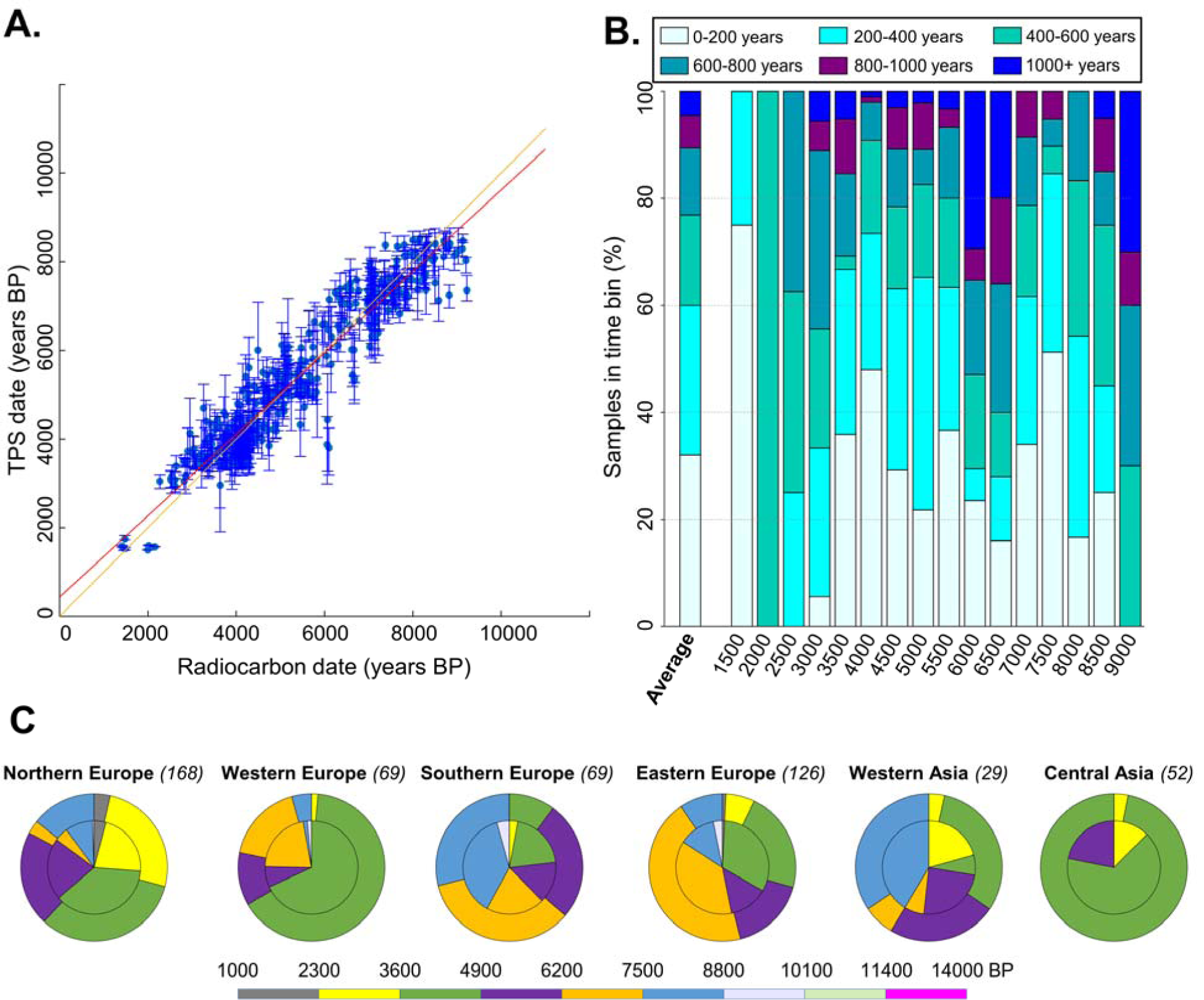
TPS results for 496 radiocarbon-dated Eurasian samples of the reference panel calculated using drop-one-individual and averaged across the 500 bootstrapping runs. A) The correlation between the radiocarbon dates and TPS results. Error bars correspond to 95% error interval. Red line represents the linear fit against the y = x line (dashed black). B) TPS aggregated accuracy. Individuals are sorted into 500-year time period bins according to their radiocarbon dates (x-axis) (e.g., the 4,000 yBP bin represents individuals dated from 4,250 to 3,750 yBP). The colours reflect the prediction accuracy, calculated as the difference in years between the mean TPS prediction and mean radiocarbon date per individual. 32% and 56% of the samples were predicted within 200 and 400 years from their radiocarbon dates, respectively. C) TPS predicted dates by region. Samples are split into 1,300 years and dated by TPS (outer pie charts) and the corresponding radiocarbon dates (inner pie charts).

To further evaluate the performances and characteristics of TPS, we applied it to six cohorts with a total of 1024 ancient and 2117 modern individuals, which were not used in its construction and are not part of its reference panel.

First, we applied TPS to 232 radiocarbon-dated Central Asians (Table S5) using the reference panel of 496 individuals yielded a median 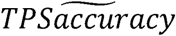 of 695 years (Table S6). The existence of reference panel samples of the same-country (*N*_*match country*_=137, 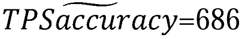, 95% CI [621, 920]; *N*_*unmatch country*_ =95, 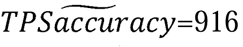, 95% CI [889, 1198]) (Fig S14) or same-country and -period as the test samples (*N*_*match country and period*_=63, 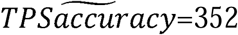, 95% CI [378, 677]; *N*_*unmatch country and period*_=169, 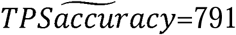, 95% CI [880, 1149]) has significantly increased the mean accuracy by 26% and 48%, respectively (two-tailed Kruskal-Wallis test, Chi-square=15.3 and 32.66, *p*=9.15*10^−5^ and 1.09*10^−8^, respectively).

Using the same reference panel, we next TPS-dated the 359 remaining Eurasians (*D*_*OD*_ and *D*_*UD*_; 16 previosuly undated) and 170 archaeologically-dated Central Asians (Table S7), which yielded median prediction accuracies of 645 ([*N*_*match country*_=307, 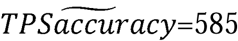, 95% CI [739, 937]; *N*_*unmatch country*_ =36, 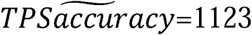, 95% CI [1158, 2426]] or same-country and -period as the test samples [*N*_*match country and period*_=269, 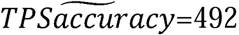, 95% CI [561, 708]; *N*_*unmatch country*_ *and period* =74, 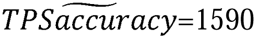, 95% CI [1683, 2400]]) and 1026 years ([*N*_*match country*_=61, 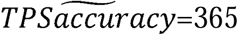, 95% CI [331, 466]; *N*_*unmatch country*_ =109, 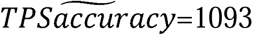, 95% CI [987, 1129]] or same-country and -period as the test samples [*N*_*match country and period*_=44, 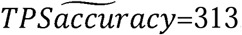, 95% CI [255, 369]; *N*_*unmatch country and period*_ =122, 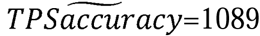, 95% CI [954, 1089]]), respectively. The existence of geographical and/or temporal reference panel samples matching to the test samples was associated with higher accuracy, particularly among Central Asians that were predicted with 12% higher accuracy than Eureaisn samples on average, suggesting the existence of geographically localized TPS patterns in Central Asia.

To test the effect of SNP numbers and using TIMs versus TPS SNPs, we next applied TPS to 121 radiocarbon-dated Europeans (Table S8) using an extended reference panel of 496 individuals as well as 443 Eurasians and Central Asian whose TPS date was less than 500 years off. Our results showed that 3,000 TIMs suffice to produce accurate dating 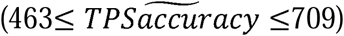 when geographical and/or temporal reference panel samples exist and that in such cases, both TIMs and using over 15,000 TPS SNPs yield comparable results (Table S9; Figs. S15-16).

To demonstrate the effects of different reference panel, we next compare the TPS results of 65 radiocarobn dated and 10 archaeologically dated Caucasus individuals (Tables S10-11). In evaluating the results against the three reference panels used thus far (Table S12), we found that while TPS accuracy was consistent with previous findings. The optimal results were obtained with the second panel consisting of Eurasians, although it was contained in the third panel, but the difference was insigificnat (two-tailed Kruskal-Wallis test, Chi-square=74, *p*=0.48). These results demonstrate a potential concern of over-saturating the reference panel.

Next, we TPS dated 67 relatives from 31 Eurasian and Central Asian families originally dated with mixed methods to improve the dating accuracy and conserve resources (Table S13). The mean date difference among families with sole or mixed radiocarbon-dated members was 294 years (*N*=50, 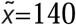, 95% CI [157, 431]), inconsistent with the observed kinship, compared to 186 (*N*=50, 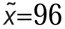, 95% CI [105, 267]) years with TPS dating. For example, the two parent-child pairs I5236-I5241 and I1378-I1732 whose radiocarbon dates were 858 and 1188 years apart were TPS-dated to overlapping periods, with the pairs being only 21 and 96 years apart, respectively. Overall, TPS dating differences of family members are smaller and more narrowly distributed compared to radiocarbon dating, but the difference is insignificant (Fig. S17). For families dated using only archaeological methods, TPS yielded a mean distance of 208 years (*N*=19, 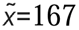, 95% CI [27, 389]).

Finally, we TPS dated modern individuals from 20 populations using the reference panel of 496 individuals supplemented by random 160 modern individuals (10 individuals per population, four populations per continent) (Table S14). TPS results (*N*=2117, 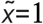, 95% CI [6, 7]) (Table S14) were consistent with their modern origins.

### TPS application to four case studies

#### 1) Dating the 13 Brandysek individuals, Czechia

Two individuals from (RISE568 and RISE569), the Brandysek site, were originally attributed by archaeological context to the Bell Beaker period (4,800-3,800 yBP) ^*17*^. Olalde et al. ^*18*^ later dated RISE569 (1,290-1,180 yBP) and RISE568 (1,350-1,150 yBP) based on the radiocarbon and archaeologically associated materials, respectively. As these post-dated the Bell Beaker Culture, the authors have omitted the two individuals from their analysis of the Brandysek individuals (4,850-4,150 yBP). TPS dated RISE569 to 3327 yBP, RISE568 to 3952 yBP, and the remaining Brandysek individuals (excepting I7272) to 3700-4117 yBP.

Three observations are noteworthy for these individuals. First, TPS’s date for RISE568 contradicts the archaeologically-derived and much younger date obtained by Olalde et al. ^*18*^ and confirms the Brandysek Bell Beaker identity of RISE568. Secondly, TPS’s date for RISE569 confirms Olalde et al.’s ^*18*^ assessment that this individual predates the Bell Beaker period by at least ∼2,000 years. Third, of the remaining eleven individuals, ten were TPS-dated to fit within the Bell Beaker and Corded Ware period (4,850-4,150 yBP). Individual I7272 was much older (5686 yBP) and predated the Corded Ware culture as is evidenced by two additional features: Firstly, I7272 lacked the ancient temporal component present in all other individuals at this site, which is ubiquitous among Bell Beaker samples and associated with the time period following the Yamnaya invasion (ancient temporal component 2) (Table S4). Secondly, I7272 Y haplogroup is I2, whereas all the other males at that site, including the other two attributed to Corded Ware, are R1. Haplogroup R1 dominates post-Yamnaya migration populations ^*27-29*^, while I2 is primarily associated with Palaeolithic and Neolithic Europe ^*30,31*^. TPS dating, temporal components, and Y-haplogroup all suggest that I7272 is related to an earlier Neolithic occupation at this site. We also note that the site consists of architectural features that are not usually associated with Bell Beaker burials, like the use of stone in graves ^*18*^. Such indefinite archaeological materials should not be used for dating.

#### 2) Establishing the Neolithic identity of Oxie 7 sample

Oxie 7 is an established Swedish Neolithic site dated to 4,900 yBP ^32^, which is often used as a type-site for the Neolithic ^33^. In their analysis of Neolithic individuals, Allentoft et al. ^17^ employed archaeological contexts to date RISE174 to the Iron Age (1,523-1,340 yBP) and excluded this individual from their analysis. TPS-date for RISE174 is 3940 yBP, consistent with the Neolithic attribution of the site.

#### 3) Correcting the radiocarbon date at Kyndeløse, Denmark

RISE61 was radiocarbon-dated to 4,801-4,442 yBP ^*17*^, which is suggestive of a Middle Neolithic origin ascribed to 5,400-4,700 yBP in Scandinavia. However, Allentoft et al. ^*17*^ also cautioned that there might be a marine reservoir effect since the ^13^C/^14^N stable isotope data were consistent with a heavily marine diet, which would potentially shift the carbon age older than a terrestrial diet by several hundred years. TPS dated RISE61 to the Late Neolithic (4186 yBP), consistent with the difference predicted by a marine reservoir effect on this individual. This finding suggests that the Passage Grave Culture at Kyndeløse ^*34*^, which began to appear around 5,300 yBP in Denmark ^*35*^ may have existed as late as 4,200 yBP.

#### 4) Tracing the origins of the Longobards

The Longobards appeared during Roman times as a barbarian tribe who lived north of the Danube in present-day Hungary. From there, they established themselves in the Roman province of Pannonia at the beginning of the sixth century CE. They subsequently invaded Italy in 568 CE and were eventually conquered by Charlemagne in 774 CE ^*36*^. A total of 10 samples were excavated from Pannonia at Szólád, a site considered to be of a Longobard type in terms of grave goods, location, and burial practices ^*37*^. Amorim et al. ^*37*^ suggested that the graves of nine of the samples (SZ2-SZ5, SZ11, SZ15, SZ36, SZ43, SZ45) are from the mid-sixth century after radiocarbon-dating one individual (SZ43: 1,475-1,355 yBP). We TPS-dated all nine individuals. TPS predictions did not overlap with the sixth-century Longobardian association (1,415-1,199 yBP) (Table S4). Amorim et al. assigned the tenth individual SZ1 to the older Bronze Age (3,800-2,900 yBP) ^*37*^, likely based on the archaeological context. TPS, instead, dated all the samples to similar periods as SZ43 (1552-1586 yBP), with SZ2 being slightly older (1646 yBP).

### Phenotypic traits connected to the TIMs

Some TIMs can be associated with phenotypes, such as those harbored in the HERC2, OCA2, and TYR genes involved in skin, eye, and hair pigmentation (Fig. 4A, Fig. S18A-B). At least since the Mesolithic, these traits were under selective pressure in favor of variants associated with lighter pigmentation ^38,39^. rs2269424 (G/A) is another TIM adjacent to PPT2 and EGFL8, genes associated with immunity. Strong evidence of selection has been found at this marker ^*31,40*^.

**Fig. 4.**
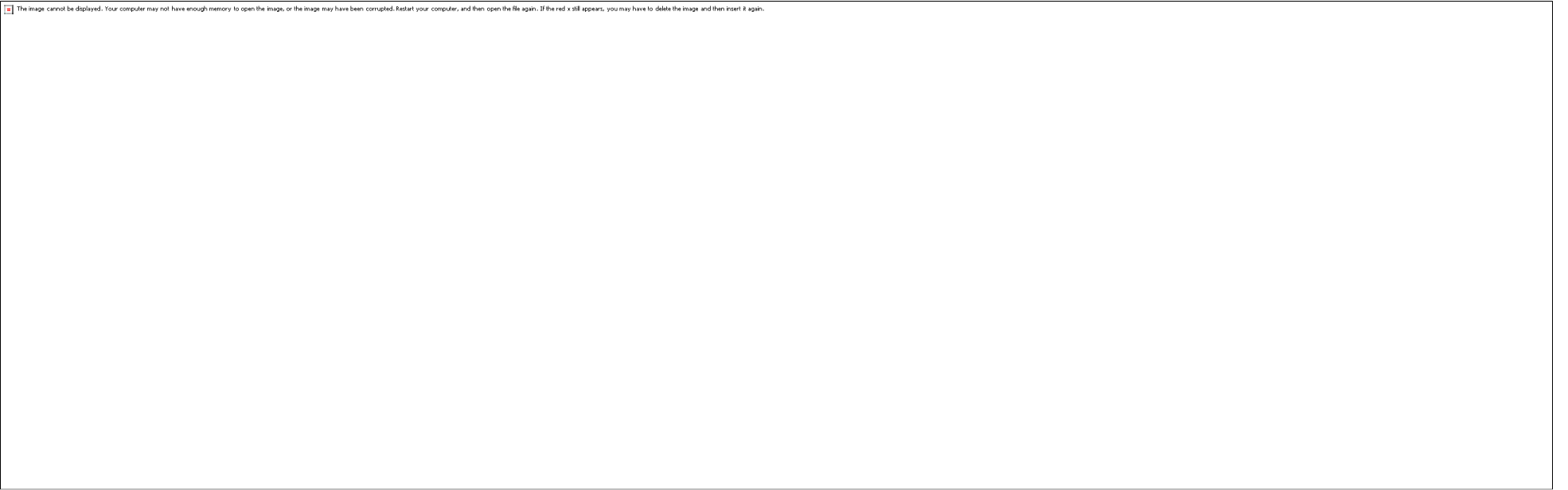
Temporal variation in the allele frequencies of three TIMs. Bars show the number of individuals genotyped for that TIM and lines show the changes in the minor allele frequency. Blue referring to radiocarbon-dated samples and green colour referring to TPS dated samples. The black line shows the weighted average of the two MAF measures. A) rs1393350 (G/A) in the TYR gene involved in pigmentation. B) rs2269424 (G/A) is adjacent to PPT2 and EGFL8 genes and associated with immunity. C) rs2073711 (A/G) in the CILP gene.

Our temporal trend (Fig. 4B) supports these findings and suggests the presence of negative selection. TIMs rs2073711 (A/G) (Fig. 4C) and rs1800562 (G/A) (Fig. S18C) located in the CILP and HFE genes are associated with cartilage scaffolding ^*41*^ and hemochromatosis ^*42*^, respectively. In both cases, the temporal variation trends indicate an increase in the frequency of these risk alleles, possibly due to the transition to a sedentary lifestyle and diet changes.

## Discussion

That about half of the ancient skeletal samples are imprecisely dated with no alternative to radiocarbon dating is a challenging problem in paleogenomics. Radiocarbon may be widely accepted as the benchmark standard for dating ancient remains ^*7,43*^, but its reliance on large amounts of organic material renders many samples undatable.

Radiocarbon dating is exposed to various environmental biases ^*20*^. By contrast, genomic dating relies on aDNA sequences, which makes it possible to directly date skeletons whose radiocarbon dating cannot be established. Motivated by our observations that allele frequencies show temporal variability over time, we introduced Time Informative Markers (TIMs) and showed their usefulness to genomic dating. For example, we showed that TIMs, like rs1393350 associated with pigmentation, have increased or decreased their frequency over time as reported elsewhere ^*31,40,42,44*^ and can be used as biomarkers for specific periods. The Temporal Population Structure (TPS) tool utilises TIMs to date ancient skeletons as far back as the Late Upper Palaeolithic using only their genomic data without prior assumptions. TPS models genomes as consisting of eight temporal components and calculates the dates by comparing the proportion of these components to those of radiocarbon-dated reference samples that fit the model. We demonstrated the accuracy of TPS by showing its ability to correctly predict over three thousand radiocarbon-dated individuals, modern individuals, ancient family relatives, and individuals with unreliable or missing dates. We further demonstrated its ability to resolve conflicting findings in the literature and increase the power of association studies by “rescuing” ancient samples.

TPS is a novel instrument to the growing toolkit of paleogeneticists that can be used when dating is unavailable or in doubt and to address contradictory findings in the Paleogenomic literature. The advantage of adopting an admixture-scheme to derive the temporal components over alternative techniques like linear regression is its resilience to missing data, common to aDNA data. Another advantage is the dating function, which weighs the ages of the temporally most-similar reference populations rather than using the closest individual, thus reducing the effects of outliers (Fig. S5). TPS’s performances depend on the number and type of SNPs included in the analysis and the comprehensiveness of the preference panel. However, over-saturating the reference panel can reduce its accuracy. As with radiocarbon dating ^2^, TPS is best used with standards and the most suitable reference panel. TPS can be used to detect outliers or misattributed samples and to develop alternative hypotheses to other dating tools. We envision that, with the increase in the number of sequenced populations over time, direct dating methods will become more accurate for broader worldwide communities. Therefore, our results should be considered a lower bound to the full potential of TPS for bio-dating. TPS is not comparable to the genetic dating method proposed by Moorjani et al. ^*21*^, which is based on the inbreeding with Neanderthal. TPS has several limitations. First, it is applicable to samples with at least 15,000 TPS SNPs or 3000 TIMs. If fewer SNPs are available, TPS predictions may be biased. Second, since TPS works best for geographically and temporally matched individuals, it cannot be used for older or non-Eurasian samples due to the sparsity of such radiocarbon-dated samples. This limitation can be addressed when more data will become available.

## Material and Methods

### Curating ancient genomic databases

We curated a database *DB*_*Anc*_ of 961 ancient individuals ^17,31,42,45-58^ and 147,229 SNPs after filtering a published ancient DNA database with 150,278 SNPs ^*24*^ with the least missingness and enforcing the following criteria: i) missingness lower than 0.9 (i.e., individuals with less than 15k SNPs were removed); ii) dates in the range 14,000-1,000 yBP; and iii) burial coordinates within Eurasia. 49 Eurasians were relatives (Data S1). We also obtained full annotation for these individuals, such as burial locations and dates (Table S1). To validate TPS we curated four additional datasets (we used the pipeline descirbed in Esposito et al. ^59^ whenever the data were not in PLINK or EIGENSTRAT format): The first included 269 radiocarbon- and 247 archaeologically-dated Central Asians, 232 and 170 of whom had over 15,000 TPS SNPs ^60^, respectively (Table S5). 18 Central Asians were relatives. The second Euroasian dataset included 143 radiocarbon-dated individuals curated from the literature (Table S6), 121 of whom were dated to 14,000 yBP ^42,48,49,54-57,61-78^ (Table S8). The third dataset included 110 mixed-dated Caucasus individuals ^79^, 75 of whom had over 15,000 TPS SNPs (Table 10). Finally, we obtained 2504 individuals from the 1000 Genomes database ^*80*^ and analysed the 2117 individuals, who were used in the construction or training of TPS (Table S14). Throughout the paper, we used mean radiocarbon dates, defined as the middle point of the dating interval obtained with the samples’ annotation.

### Constructing the temporal components

To identify the temporal components, we randomly selected 300 samples from *DB*_*Anc*_ that were merged with 250 individuals (*DB*_*Mod*_) selected randomly and evenly from five present-day populations (Chinese CHB, Yoruba YRI, Finnish FIN, British GBR, and Tuscan TSI) ^*80*^, and filtered them for the TPS SNPs. We applied *unsupervised* ADMIXTURE (ver 1.3) ^*25*^ for *K’s*, ranging from 4 to 12. Individuals were sorted by time and temporal plots were produced for each run (Fig. S1). For each plot, we selected putative temporal components that were characteristic of time (not geography), i.e., components that were evenly distributed in all samples within a certain time period. Ten and three putative modern and ancient components were identified, respectively, with a clear split between the ancient and modern components. We continued refining them separately.

For the ancient components, using the allele frequencies output of ADMIXTURE (*p*-file), 15 synthetic samples associated with each temporal component candidate were generated ^*24,81*^. A Principal Component Analysis (PCA) plot of these 150 samples showed the components to be separated, except for two that could not be resolved. We discarded one of the two and were left with 135 synthetic samples from nine components (Fig. S2). We merged these 135 synthetic samples with the 300 ancient individuals and ran ADMIXTURE in a supervised mode against the synthetic samples. In the following supervised ADMIXTURE analyses, we dropped one component at the time and compared the resulting plots, reducing the number of overlapping components. Eventually, five ancient temporal components were retained (Fig. S3; Table S2). For the modern components, we found that three components best described our samples and that when merging them with the ancient individuals, *K*=8 had minimum noise, smoother profiles and high ancient-modern sample separation (Fig. S4) (Data S2). In all the supervised analyses the gene pools were inferred in relation to the synthetic samples, which remained constant.

### Calculating the TPS reference panel

Based on their date, radiocarbon-dated individuals *DB*_*RD*_ were grouped into non-overlapping bins of 500 years ([14,250-13,750] … [1,250-750] yBP). We calculated the temporal componetns of those individuals by applying a *supervised* ADMIXTURE solely against the temporal components (Data S2). For each of 500 TPS runs, 400 samples were chosen at random with stratified sampling to represent all the temporal bins. We then applied, for these samples, an iterative combination of *k*-means (starting from *k* = 2) and pairwise *F*-tests within each group with at least three reference individuals, to identify possible substructures based on their temporal profiles. For each group and *k*, we applied *k*-means to the temporal coefficients of the reference individuals to identify the clusters and their centroids. A pairwise *F*-test between centroids was used to verify the hypothesis that each cluster was indeed a distinct entity from all the others. If this hypothesis was verified for all the clusters within the group, the next value of *k* was tested. Alternatively, the previous *k* represented the optimal number of subgroups that describe the genetic variability within the temporal bin. Each subgroup was called a temporal population, characterized by an average temporal signature and an average date *T* with the corresponding error calculated as a standard deviation of the components. Temporal populations of only one individual were discarded. The remaining temporal populations were added to the reference panel (Table S4).

### Predictive algorithm

To date a test sample, we used a *supervised* ADMIXTURE, which infers its temporal components only against the temporal populations of the reference panel (Data S2). TPS then calculates the genetic distances *{g*_*i*_*}*_*i=*1…*N*_ between the individual and the temporal populations as the Euclidean distance between two temporal vectors. Finally, TPS selects the three temporal populations *P*_*1*_, *P*_*2*_, *P*_*3*_ with the closest genetic distance to the test sample and applies a weighted average to their average dates *τ = w*_*1*_*T*_*1*_ *+ w*_*2*_*T*_*2*_ *+ w*_*3*_*T*_*3*_, where the weights (*w*_*i*_) are the inverse of the genetic distances normalized to their sum

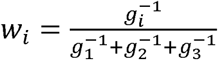

and *τ* is the TPS date for the test sample. This procedure is repeated 500 times and TPS returns the average date and the error intervals, which represent the uncertainty in the admixture results. TPS results reported here are the average of the 500 bootstrapping runs.

### Evaluating the accuracy of TPS

All analyses were done using the leave-one-out individual procedure, where each individual is tested against a reference panel reconstructed without it, unless when new datasets were tested whose samples were not used for the calibration or training of TPS. When testing parent-offspring pairs on the original cohort, the tested pair or triples were omitted from the reference panel. A tested individual *matched* the country and date of the reference panel if its country was included in the reference panel and the difference between its published date and that of reference samples from that country was less than 500 years.

### TPS application to four case studies

We evaluated the case studies against three reference panel (Table S15): 1) the standard one with 496 individuals (Table S4), 2) the standard one supplemented with Eurasian samples that TPS predicted within less than 500 years of their estimated date (Table S4) and 3) the latter dataset supplemented with Central Asian samples that TPS predicted with similar accuracy (Data S3). Since all the case studies were European and because the radiocarbon-dated samples best matched the European reference panels, we used the second reference panel for TPS dating, and note that over-saturation of the reference panel decreases its accuracy.

### Identifying Time Informative Markers (TIMs)

Provided the per-SNP allele frequencies (ADMIXTURE’s *p*-file), which comprise the five ancient temporal components and represent different periods, it was possible to associate the temporal components with time periods (Figs. S3, S6). Sorting the five ancient temporal components from old to new, we used the allele frequencies of each component (Table S3) to detect SNPs whose allele frequencies show directed behavior over time. For that, we constructed a time series with the date ranges assigned to the temporal components in 500-year bins (from 14,000 to 3,500 yBP), resulting in 22 data points (Fig. S6). Overlaps in the assigned dates range of the temporal components were averaged to construct the time series, and the resulting temporal trends were smoothed using a moving average filter to reduce noise. A total of 62,371 SNPs showing global increasing or decreasing trends or displaying local behavior over sub-intervals of at least 3,000 years were considered TIMs (Fig. S7; Table S3). 24,311 SNPs, whose allele frequencies exhibit little or no variation over time (absolute change <=0.1) were considered non-TIMs. 715 individuals with at least 15k TIMs were analyzed.

### Null models

We gauged the extent to which the choice of markers affected TPS’s accuracy by constructing two null models. The first one comprised of 24,311 non-TIMs and 385 individuals with at least 15k non-TIMs. The second one, comprised of the complete SNP-set. Here, we used random temporal components drawn from a uniform distribution.

## Supplementary information

**Fig. S1.**
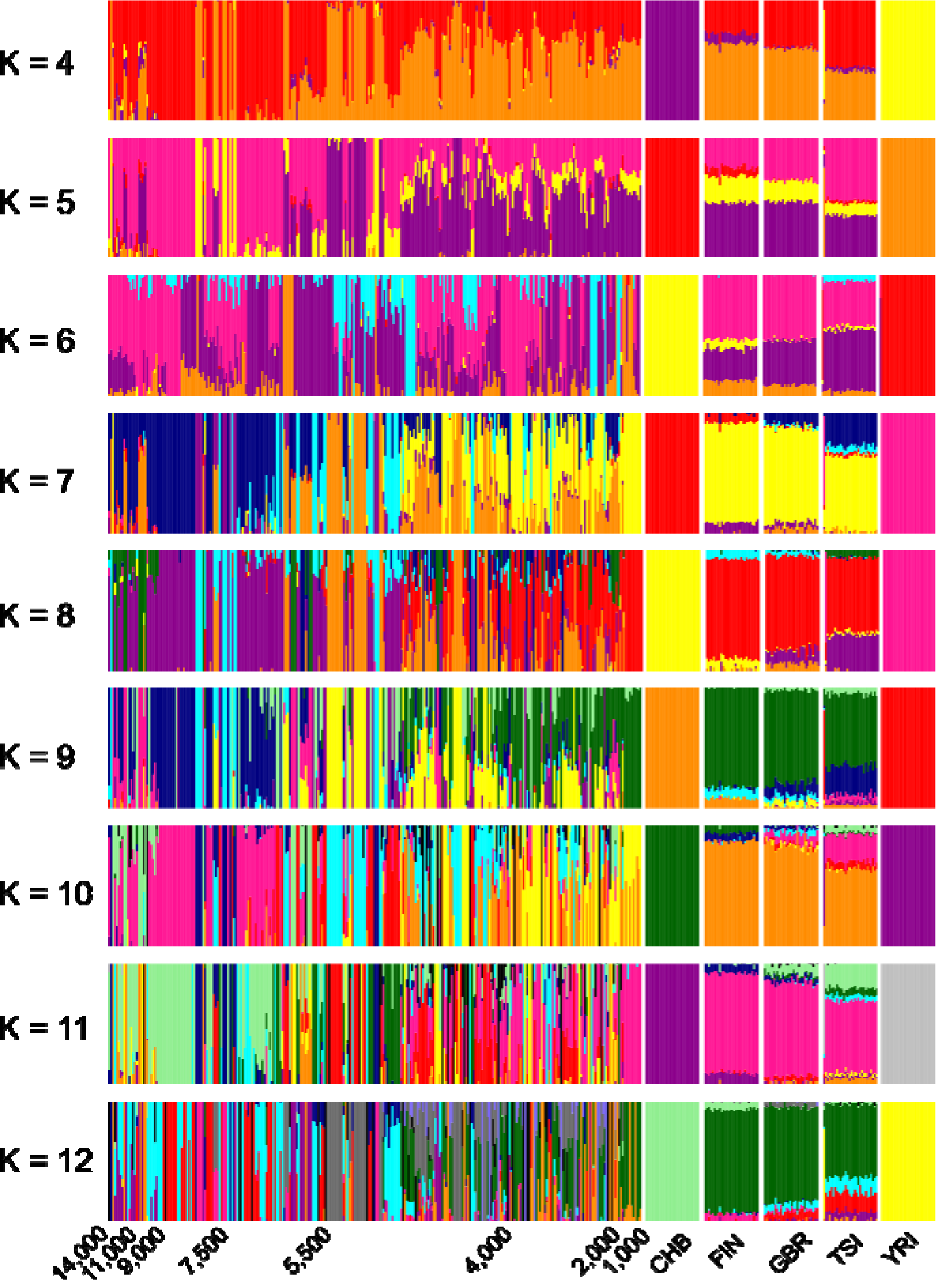
Temporal plots of 300 ancient and 250 modern 1000-genomes individuals sorted by age. The plots were obtained using *unsupervised* ADMIXTURE with *K’s*, ranging from 4 to 12. Each vertical stacked bar represents an individual. Colours correspond to the temporal components.

**Fig. S2.**
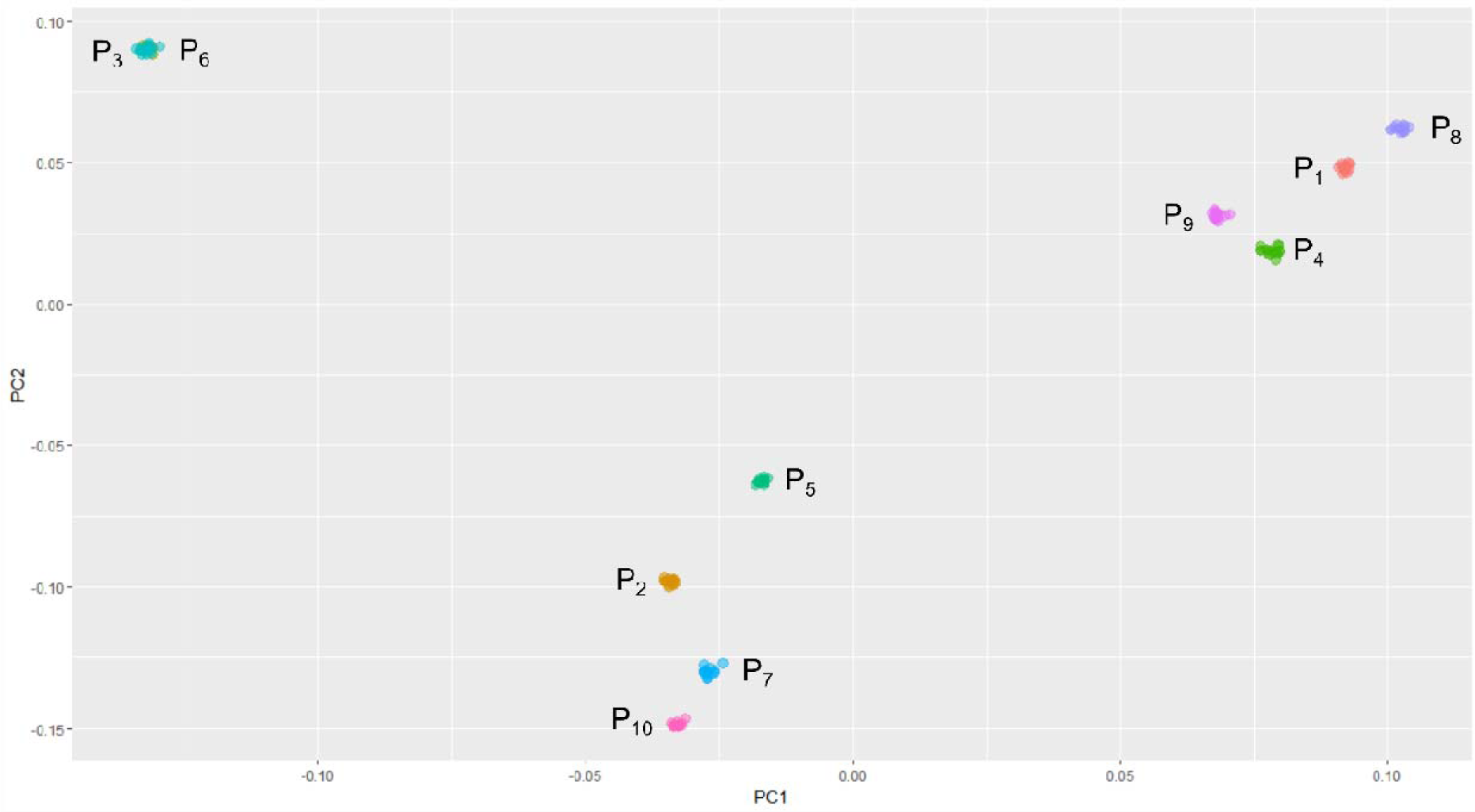
Principal Component Analysis (PCA) plot for 150 simulated samples derived from ten ancient temporal component candidates (15 samples per component). Two clusters (P_3_ and P_6_) showed a complete overlap (top left corner) and one of them was removed.

**Fig. S3.**
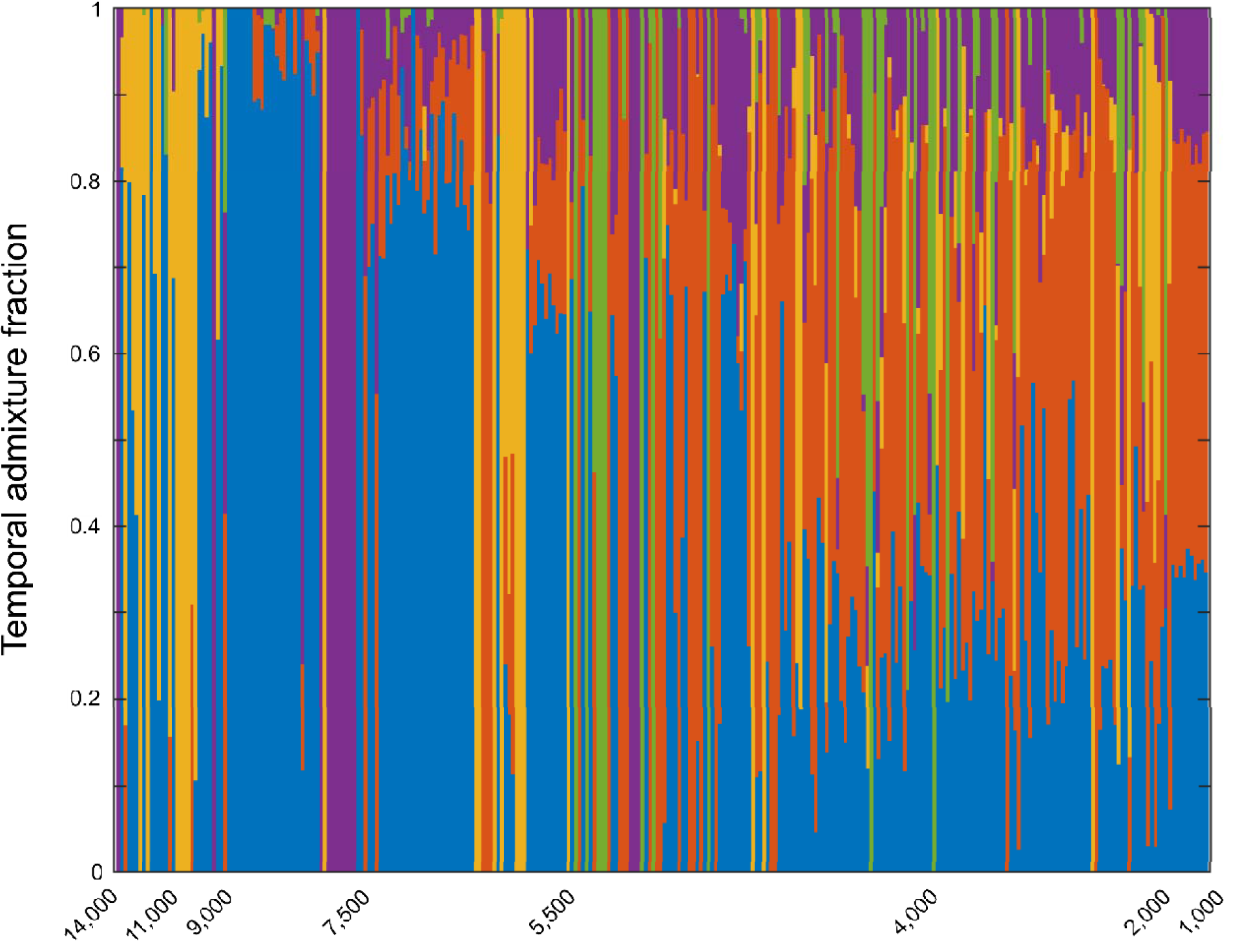
Ancient temporal components of 300 ancient individuals sorted by age. Results were obtained using *supervised* ADMIXTURE with five ancient temporal components. Each vertical stacked bar represents an individual. Colours correspond to the temporal components.

**Fig. S4.**
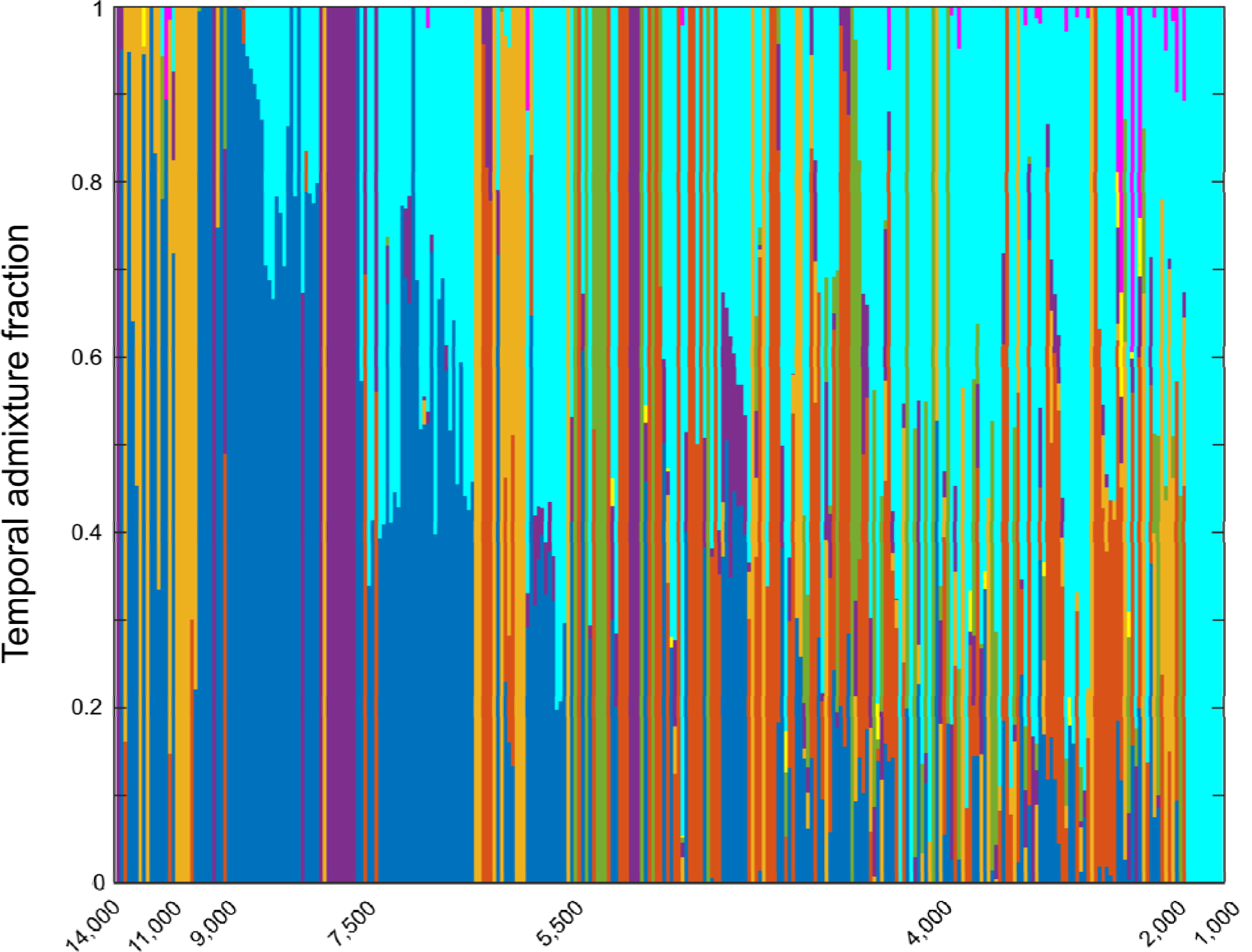
All temporal components for 300 ancient individuals sorted by age. The results were obtained using *supervised* ADMIXTURE with the five ancient and three modern temporal components. Each vertical stacked bar represents an individual. Colours correspond to the temporal components.

**Fig. S5.**
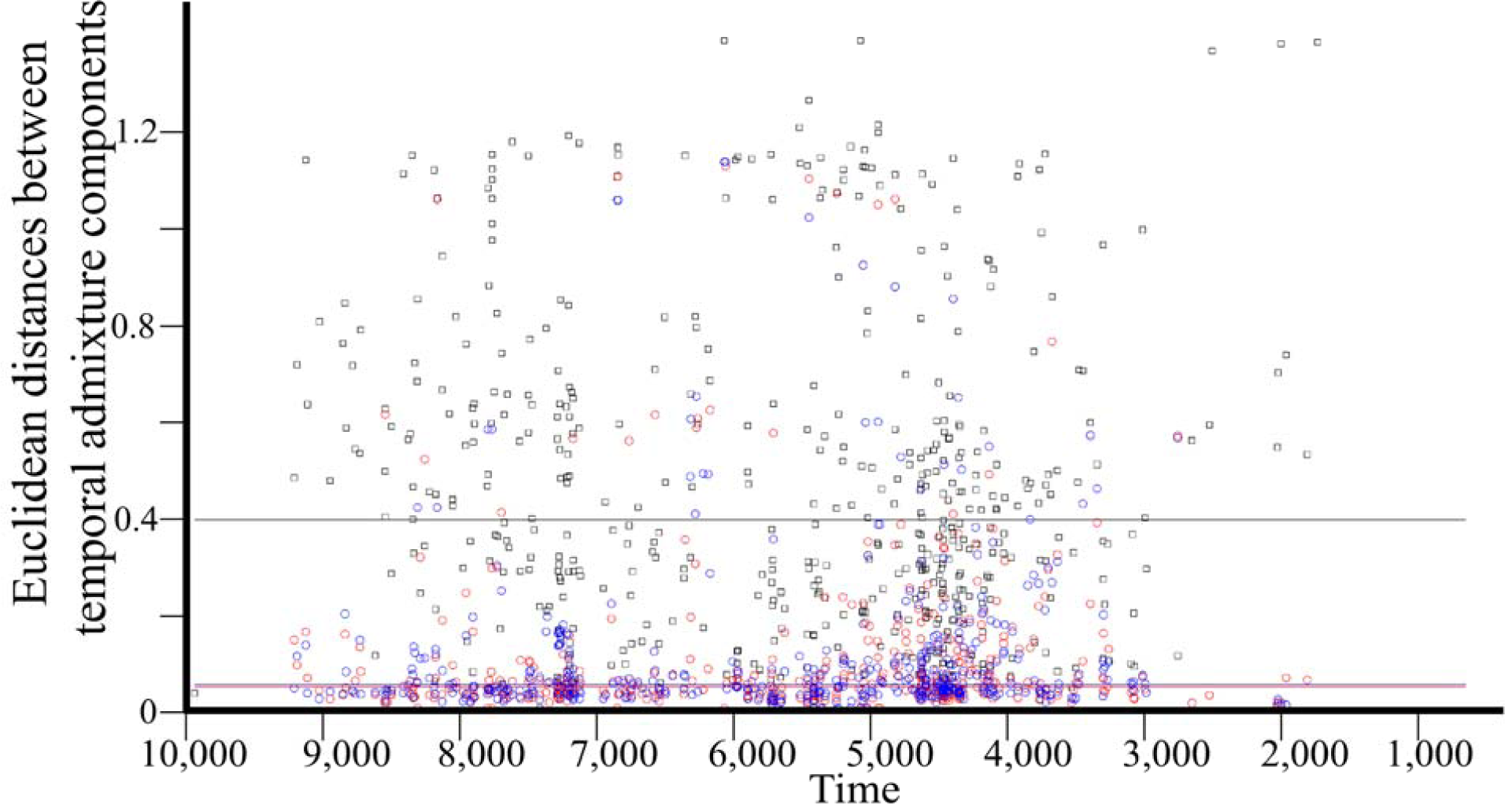
A scatter plot of the temporal distances between 496 reference panel individuals of comparative ages (Fig. 3). For each individual, we found the two closest individuals within 0-200 and 201-500 years and calculated their temporal distances from the test individual using the Euclidean distance of their eight components. We also calculated the distance to a random individual. The temporal distances were plotted: 0-200 years (red circles), 0-500 years (blue circles), and the distances to the random individuals (black squares). The median of each measure is shown as a horizontal bar (*N*_*0-200*_=492 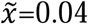;*N*_*200-500*_=490 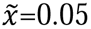; *N*_*random*_=496 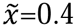). These results demonstrate the high similarity in the temporal components of individuals of similar ages. This property allows predicting ages based on the similarity between the temporal components of the test and reference individuals. We also note the existent of outliers, which requires considering multiple samples from the reference panel rather than the one with the shortest distance.

**Fig. S6.**
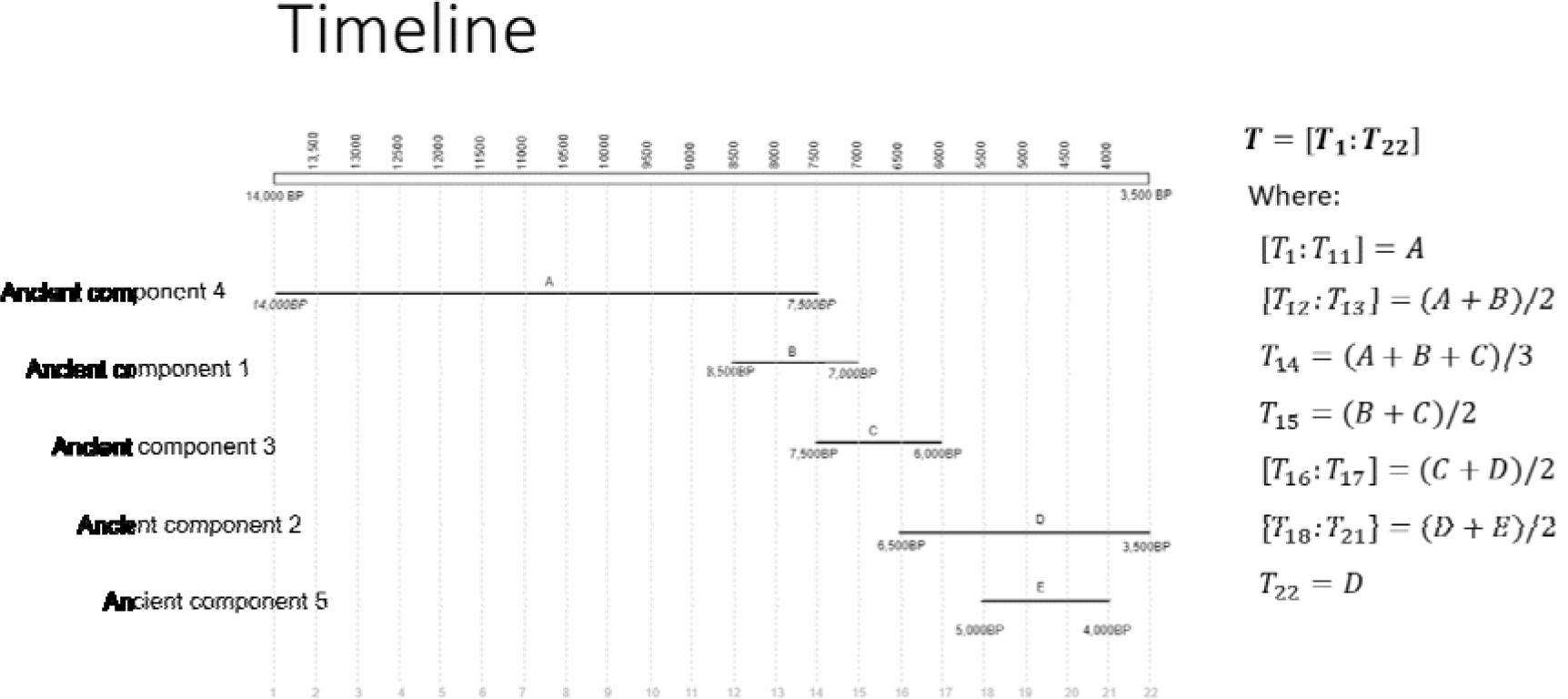
Construction of time series for the identification of Time Informative Markers (TIMs). The five ancient temporal components (Fig. S3) were assigned a time period in our timeline, as shown by the five horizontal bars. The timeline was divided into 22 bins of 500 years. For each SNP we constructed a time series *T* = [*T*_1_:*T*_22_] that consists of 22 allele frequencies (legend). Values were obtained from the allele frequencies of the five temporal components for that SNP, combined as shown in the legend. For instance, the values of *T*_1_ to *T*_11_ were equal to the allele frequency of the ancient temporal component 4 (labelled as A), as this component was dominant during this time interval; the values of *T*_12_ and *T*_13_ were the average between the allele frequency of ancient temporal components 4 and 1 (labelled as B), as both components were dominant in this time interval. The final time series were smoothed using a moving average filter to reduce noise. For example, following this procedure, TIM rs6603791 showed a frequency of 0.71 for the 10,000 yBP bin and a frequency of 0.47 for the 3,000 yBP bin, with decreasing intermediate values. The time series of the TIMs are shown in Table S3.

**Fig. S7.**
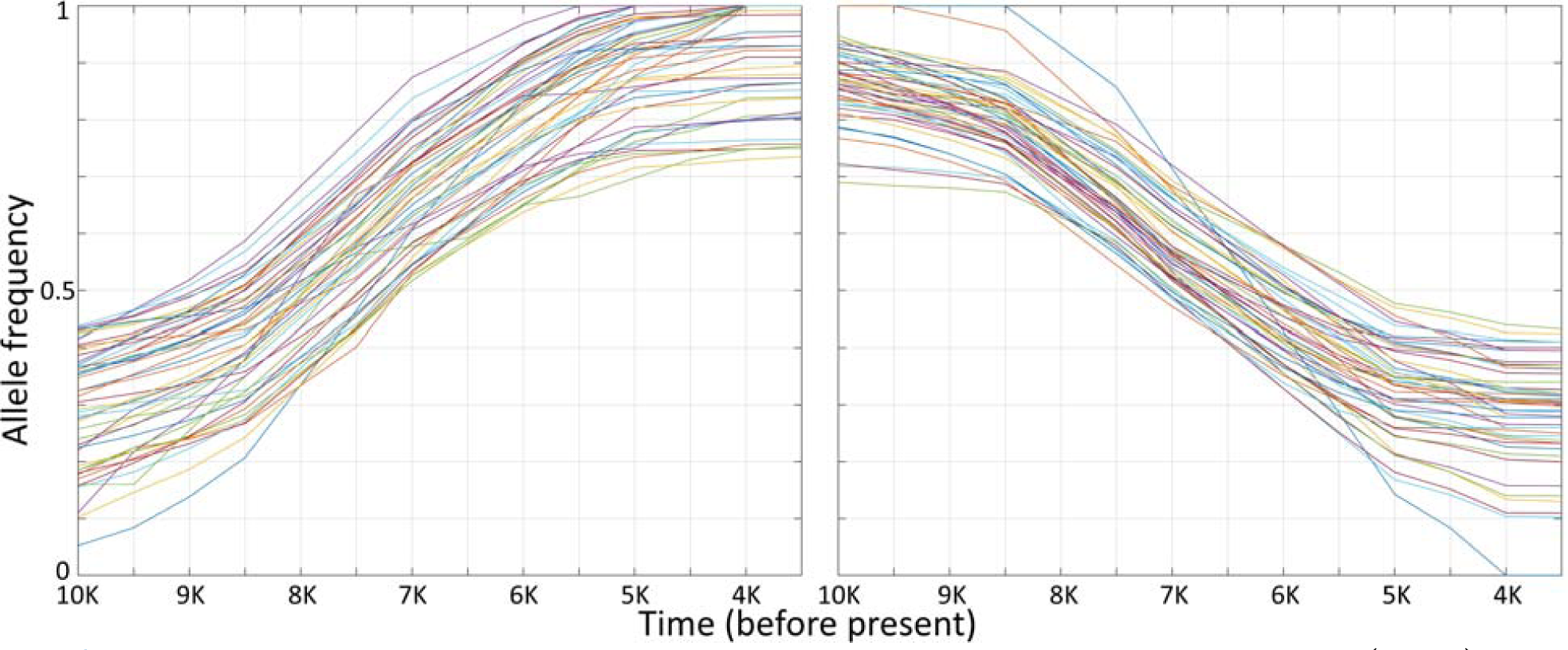
Time series of minor alleles frequencies for 100 Time Informative Markers (TIMs) that showed the most pronounced 50 increasing (left) and 50 decreasing (right) trends.

**Fig. S8.**
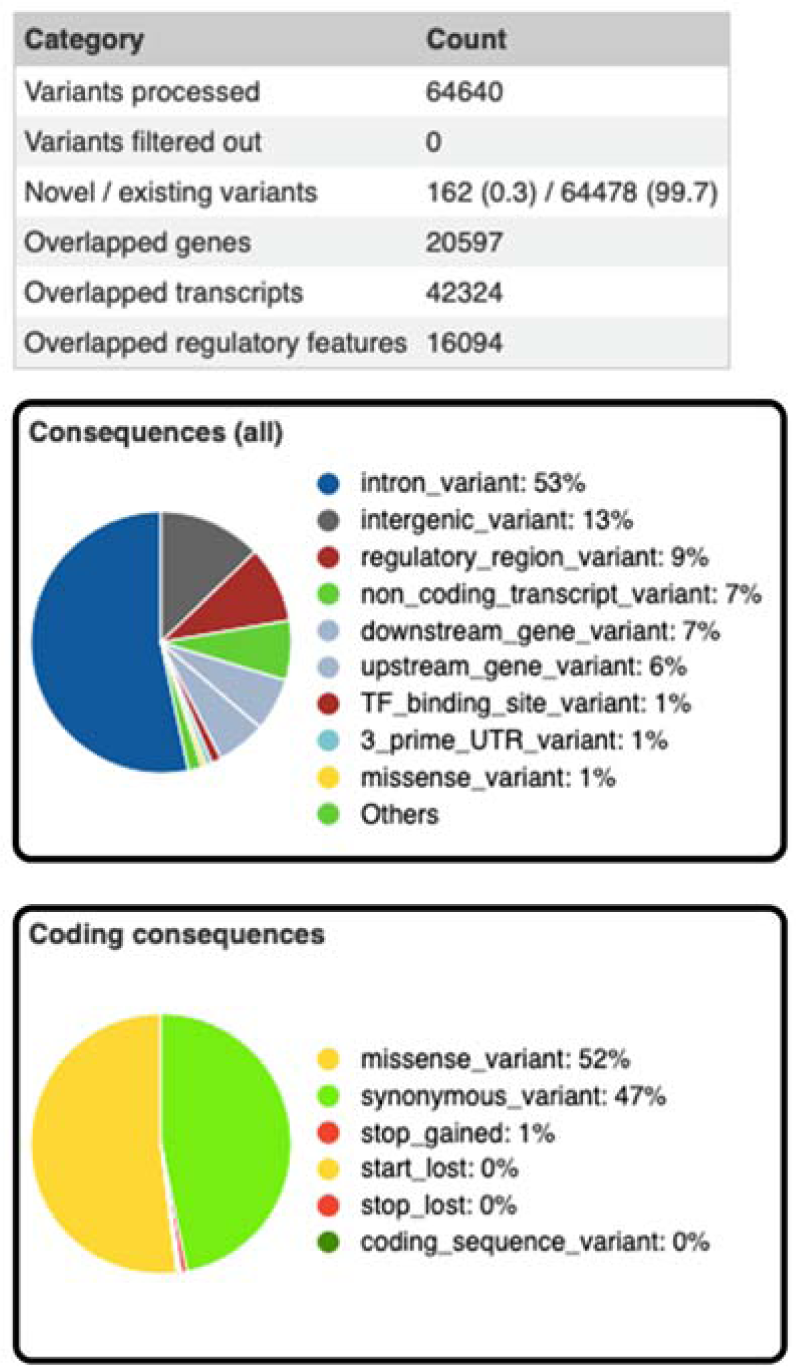
VEP summary results for the Time Informative Markers (TIMs).

**Fig. S9.**
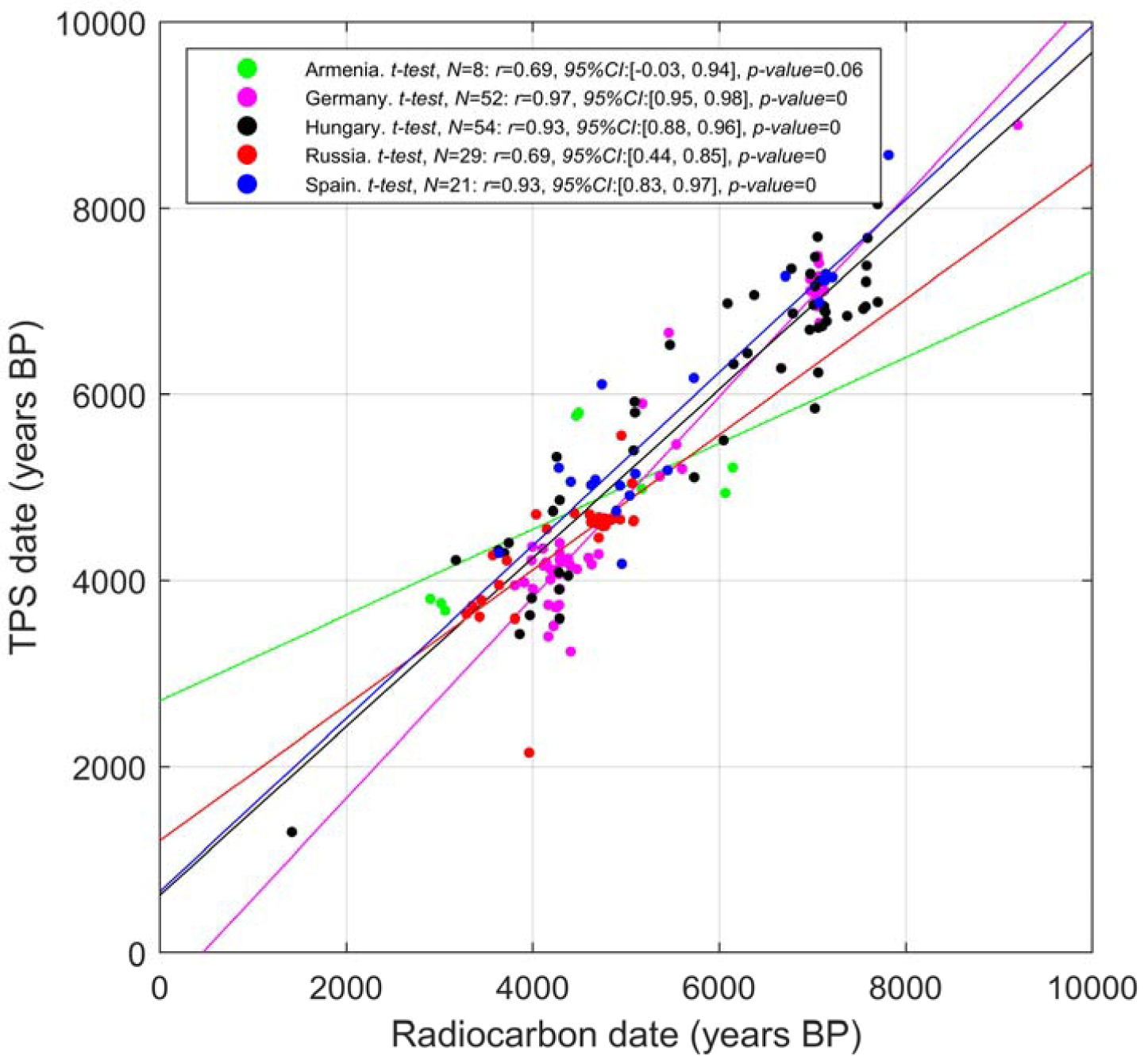
Correlation between radiocarbon dates and TPS results for 164 same-country radiocarbon-dated individuals (from Fig. 3A) averaged across the 500 bootstrapping runs. TPS results were distributed along the timeline that follows their radiocarbon dates, confirming that the temporal components represent temporal rather than a geographical variation. The only countries diverging from this general behaviour are Armenia and Iran, suggesting the possibility that some geographic effect might be present in their components (Table S4).

**Fig. S10.**
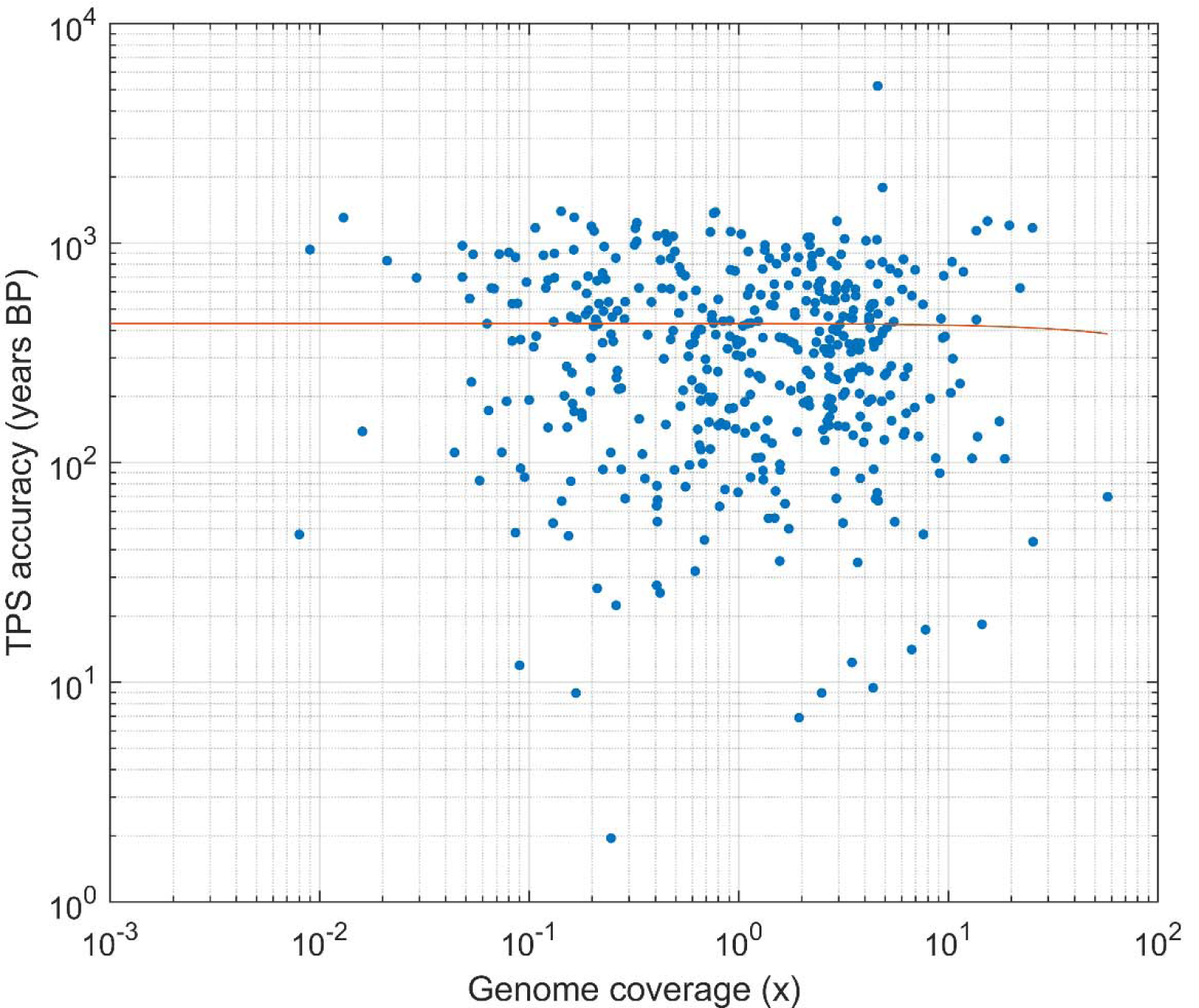
Correlation between TPS accuracy and genome coverage for the radiocarbon-dated samples for which coverage was reported in the literature. Both axes are logarithmically scaled, the red line represents the linear fit. There is no correlation between the two quantities (two-sided *t-test, N*=496: *r*=-0.0085, *95% CI*: [-0.1026, 0.0858], *p*=0.8607), suggesting that TPS results are not sensitive to missingness (provided the use of at least 15k SNPs).

**Fig. S11.**
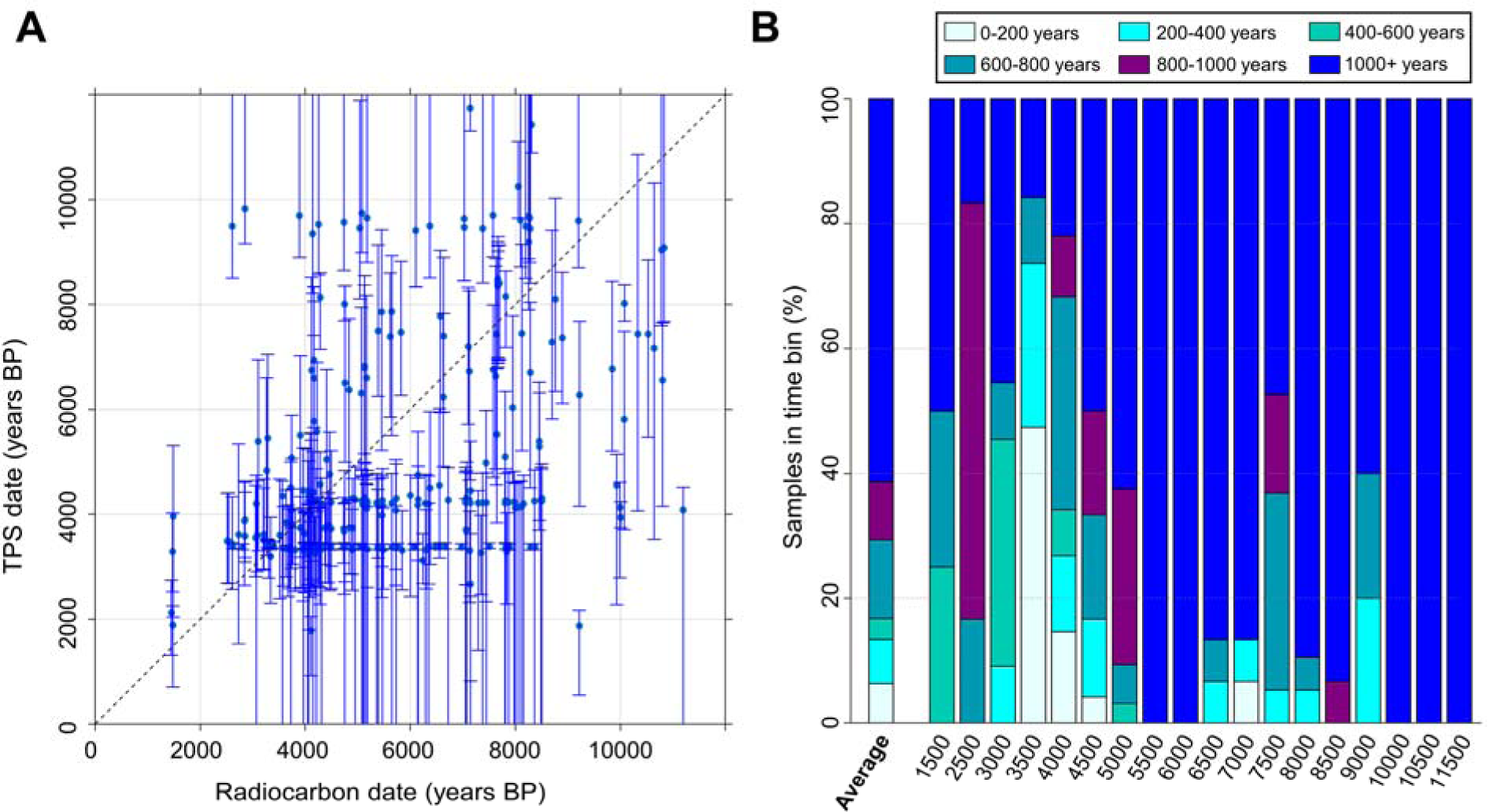
TPS results for 276 radiocarbon-dated Eurasian samples with the 24,311 non-TIMs, averaged across 500 bootstrapping runs. A) The correlation between radiocarbon dates and TPS results for the testing dataset (two-sided *t-test, N=*276: *r*=0.39, *95% CI*: [0.282, 0.486], *p*=4.1e-11). Error bars correspond to a 95% error interval. The dashed black line represents the *y = x* line. B) TPS aggregated accuracy. Individuals are sorted into 500-year bin periods according to their radiocarbon dates (*x*-axis). The colours reflect the prediction accuracy, calculated as the difference in years between average TPS prediction and mean radiocarbon date per individual. Note that due to the higher missingness in the data that resulted from the smaller SNP set, the non-TIMs dataset required dropping an additional 219 individuals compared to the all-SNP dataset.

**Fig. S12.**
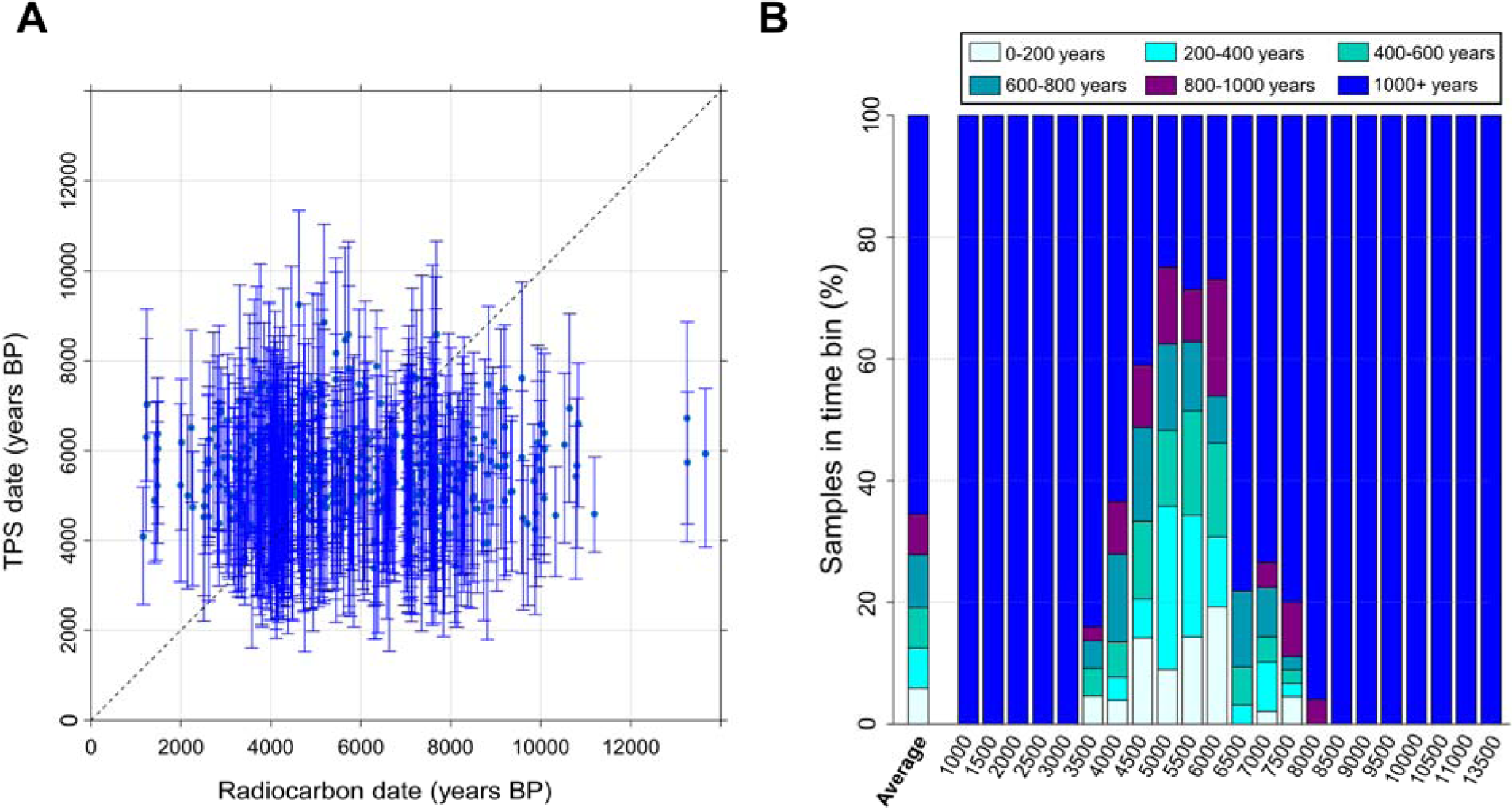
TPS results for 602 radiocarbon-dated Eurasian samples with a random model averaged across 500 bootstrapping runs. A) The correlation between radiocarbon dates and TPS results for the testing dataset (two-sided *t-test, N*=602: *r*=0.04, *95% CI*: [-0.038, 0.122], *p*=0.295). Error bars correspond to a 95% error interval. The dashed black line represents the ideal *y = x*. B) TPS aggregated accuracy (*x*-axis). The colours reflect the prediction accuracy, calculated as the difference in years between average TPS prediction and mean radiocarbon date per individual. Individuals are sorted into 500-year bin periods according to their radiocarbon dates.

**Fig. S13.**
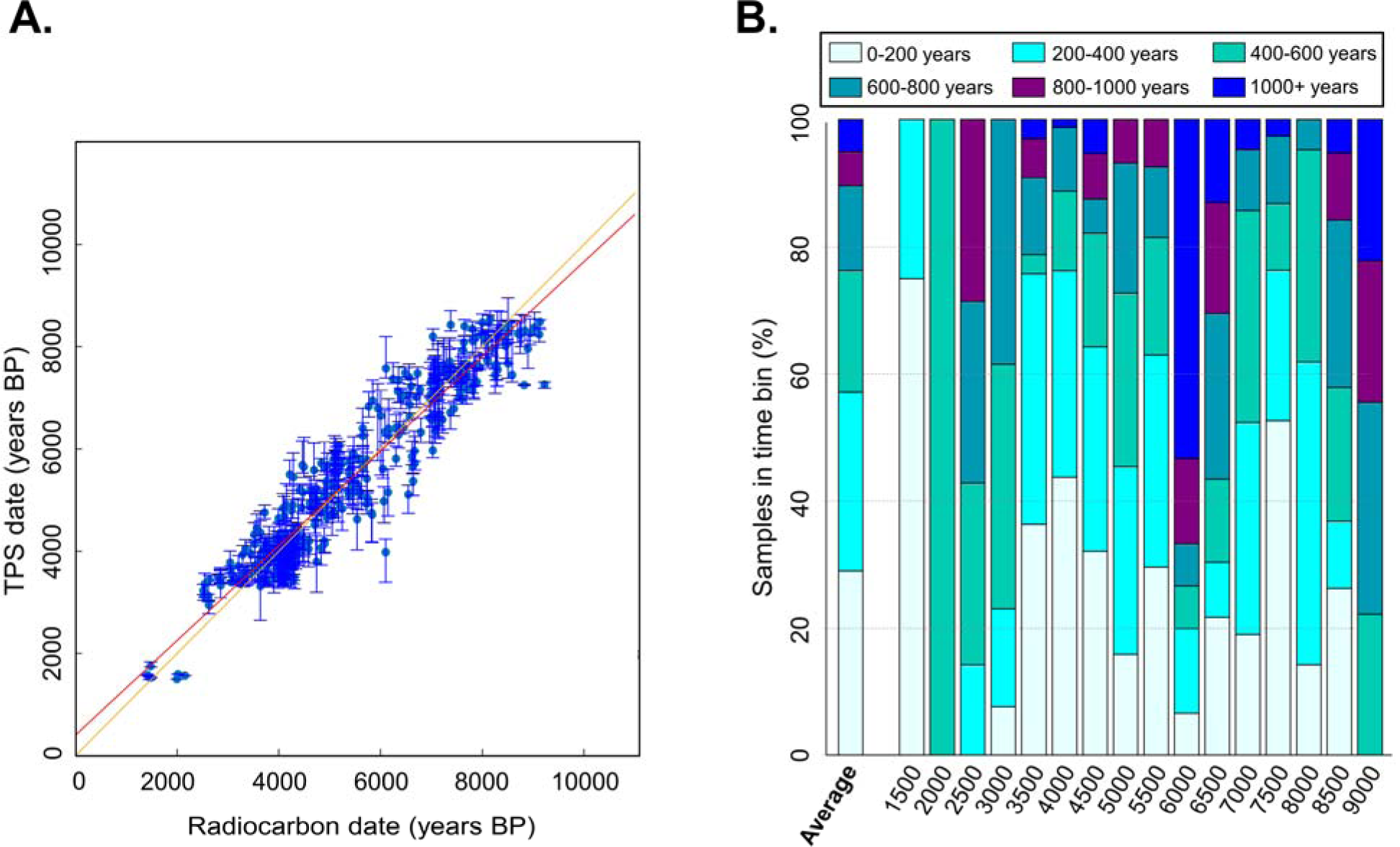
TPS results for the 437 radiocarbon-dated Eurasian samples with TIMs calculated with the drop-one-individual test averaged across the 500 bootstrapping runs. A) The correlation between radiocarbon dates and TPS results for the testing dataset. Error bars correspond to a 95% error interval. The red line represents the linear fit against the *y = x* line (dashed black). B) TPS aggregated accuracy. Individuals are sorted into 500-year bin periods according to their radiocarbon dates (*x*-axis). The colours reflect the prediction accuracy, calculated as the difference in years between average TPS prediction and mean radiocarbon date per individual. These results (*t-test, N*=437: *r*=0.96, *95% CI*: [0.948, 0.964], *p*=1.2e-215) replicated those obtained with the full set of markers shown in Fig. 3. Note that due to the higher the missingness, which was a result of the smaller SNP set, the TIMs dataset consisted of fewer individuals compared to the TPS SNP dataset. The total number of individuals studied in this dataset is 816, of which 521 were radiocarbon dated.

**Fig. S14.**
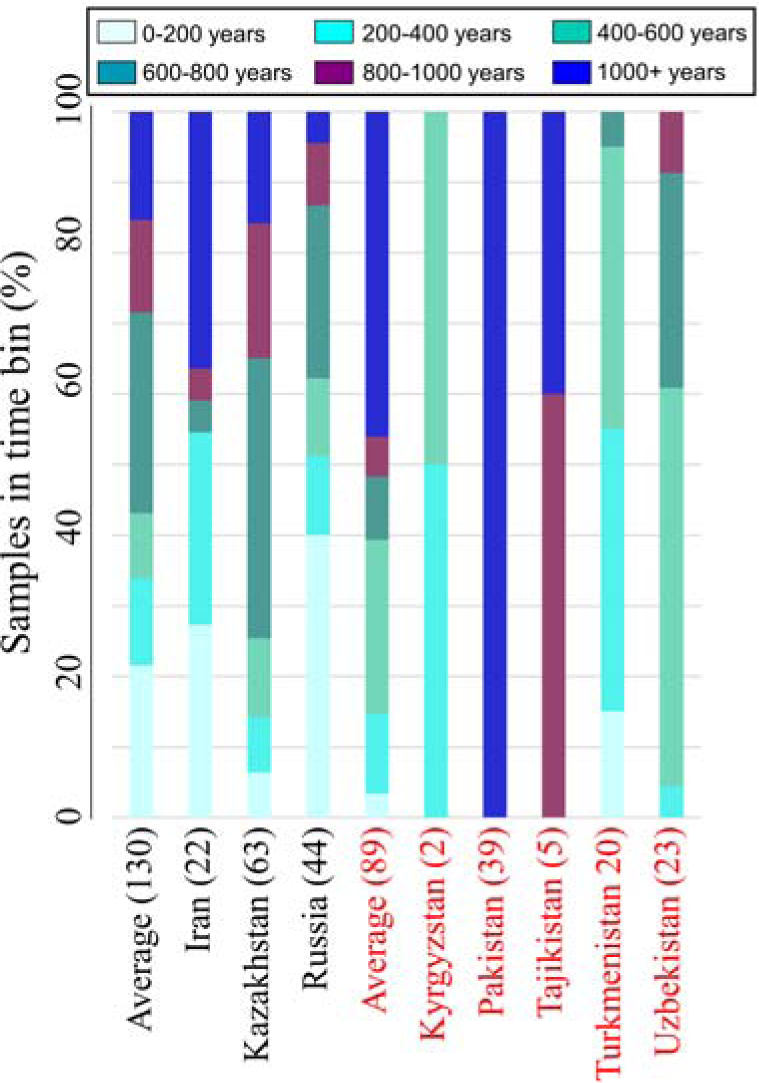
TPS aggregated accuracy for 219 radiocarbon dated Central Asians. Individuals are sorted by countries that are either represented (black) or unrepresented (red) in the reference panel. The average accuracy for each section is provided. The number of individuals per country is noted. The colours reflect the prediction accuracy, calculated as the difference in years between the mean TPS prediction and mean radiocarbon date per individual.

**Fig. S15.**
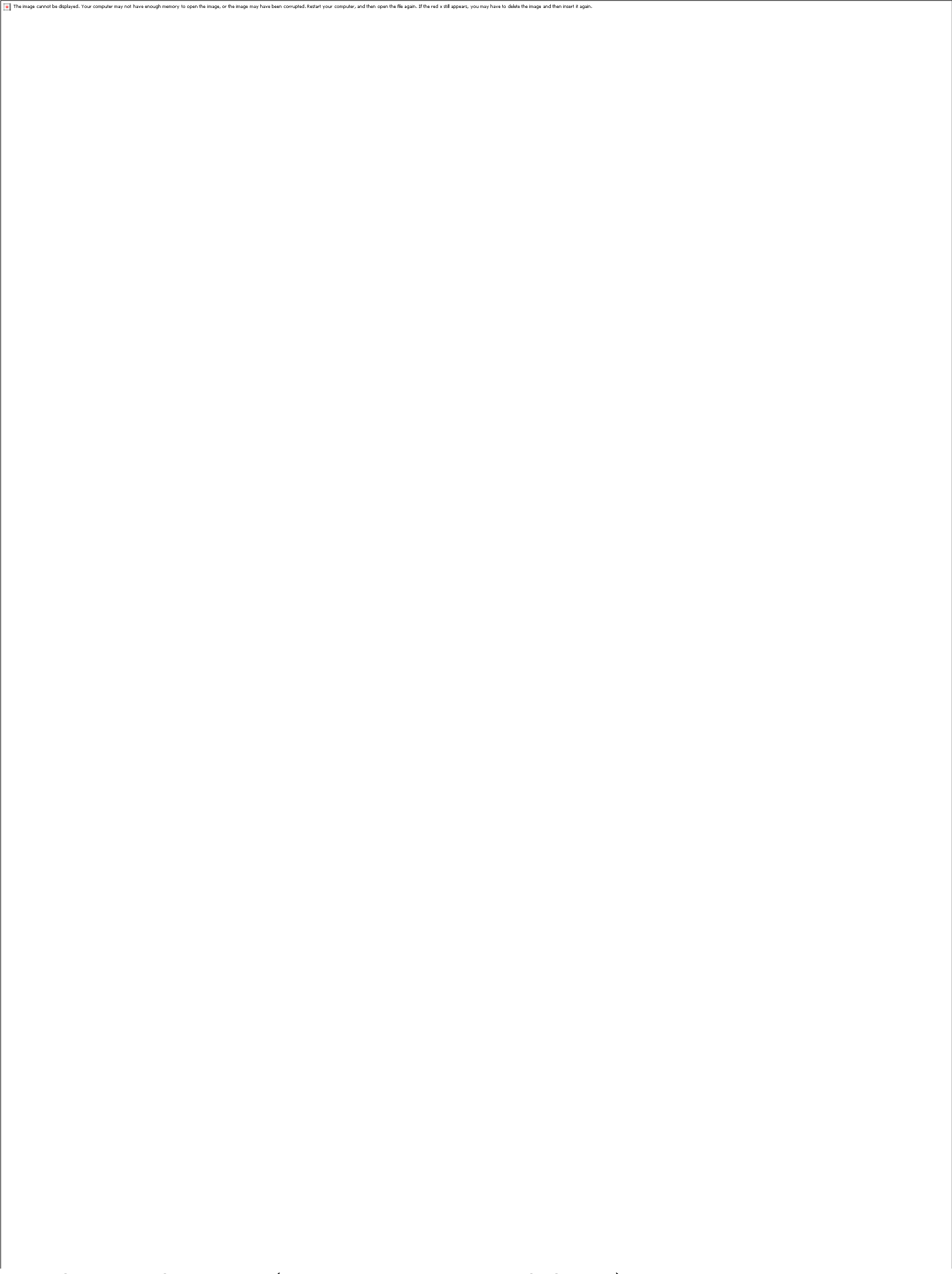
TPS results (calculated using TPS SNPs) for 121 radiocarbon-dated Europeans using various settings, averaged across the 500 bootstrapping runs. Each figure exhibits the correlation between radiocarbon dates and TPS dates for a subset of the dataset. Error bars correspond to a 95% error interval of TPS and the radiocarbon dates. The red line represents the linear fit against the *y = x* line (dashed black). TPS was applied to the samples using all the SNPs that overlapped with TPS SNPs. TPS was then run on all the samples (A-B), on test samples that have a matching reference sample in terms of country and period (C-D), and test samples that did not have a match in the reference panel (E-F). Samples in the left column (A, C, E) have over 10,000 TPS SNPs. Samples in the right column (B,D,F) have over 15,000 TPS SNPs. More details on this analysis are in Table S9 (left column).

**Fig. S16.**
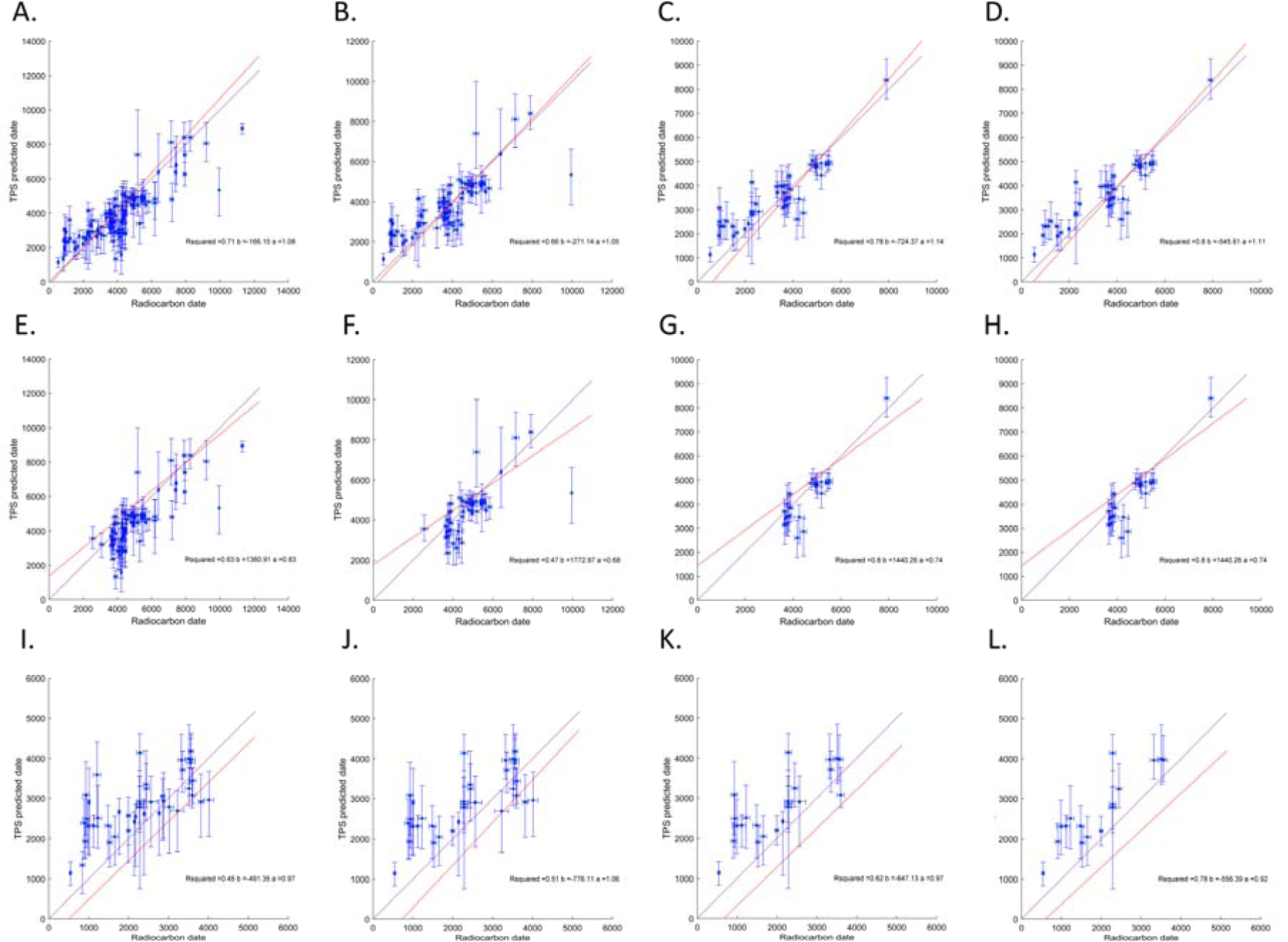
TPS results (calculated using TIMs) for 121 radiocarbon-dated Europeans using various settings, averaged across the 500 bootstrapping runs. Each figure exhibits the correlation between radiocarbon dates and TPS dates for a subset of the dataset. Error bars correspond to a 95% error interval of TPS and the radiocarbon dates. A red line represents the linear fit against the *y = x* line (dashed black). TPS was applied to the samples using all the SNPs that overlapped with the TIMs. TPS was then run on all the samples (A-D), on samples that have a matching reference sample in terms of country and period (E-H), and test samples that did not have a match in the reference panel (I-L). The four columns show samples that vary over the number of TIMs analyzed (from left to right): 3,000, 5,000, 10,000, and 15,000 TIMs. More details on this analysis are in Table S9 (right column).

**Fig. S17.**
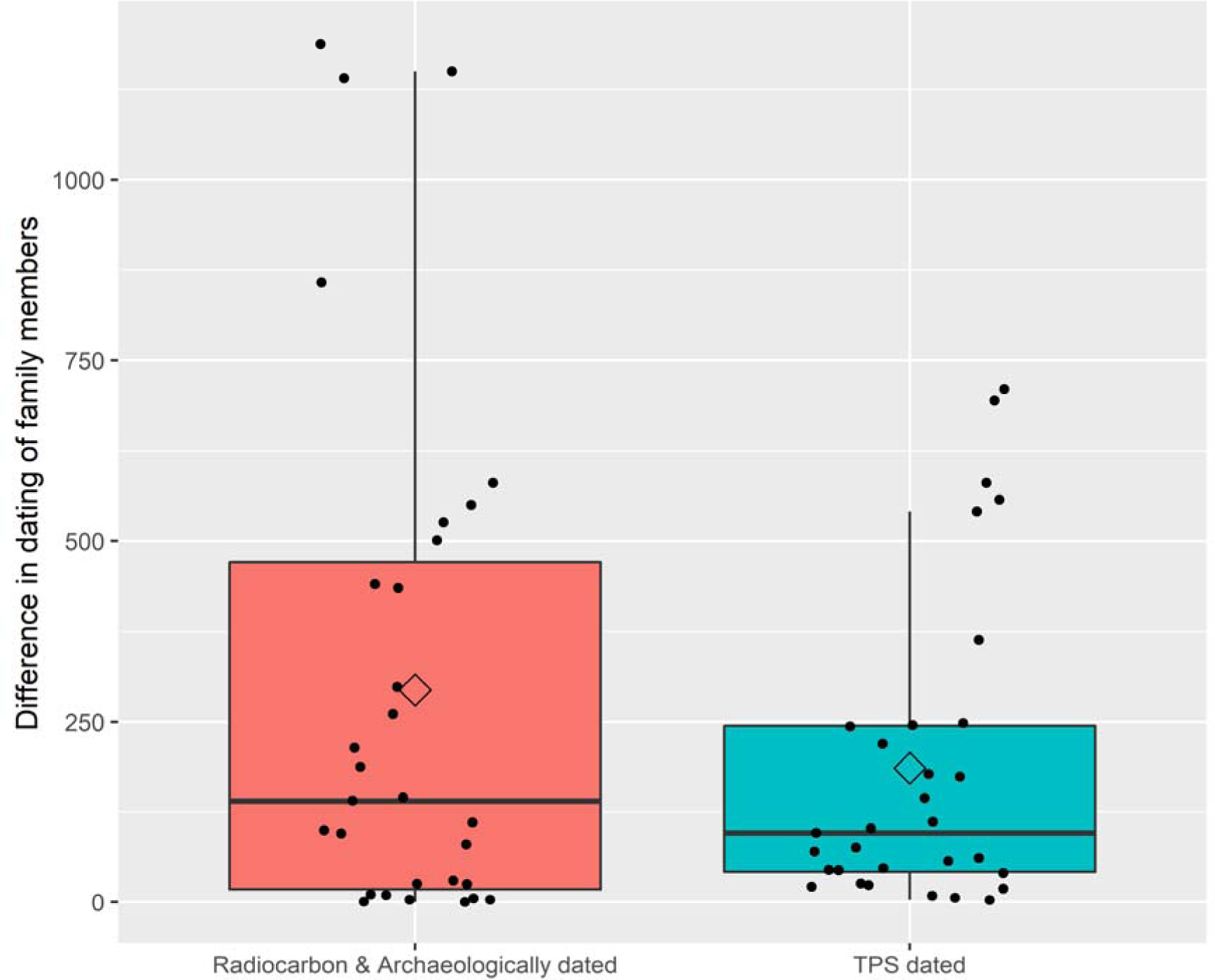
Dating differences between all family members of the same family calculated using the published dates and TPS dates. Diamonds show the mean. TPS dates are characterised in lower mean and median and narrower distribution, but the difference between the distributions is insignificant (two-sided Kruskal-Wallis test, Chi-square=0.1, *p*=0.76).

**Fig. S18.**
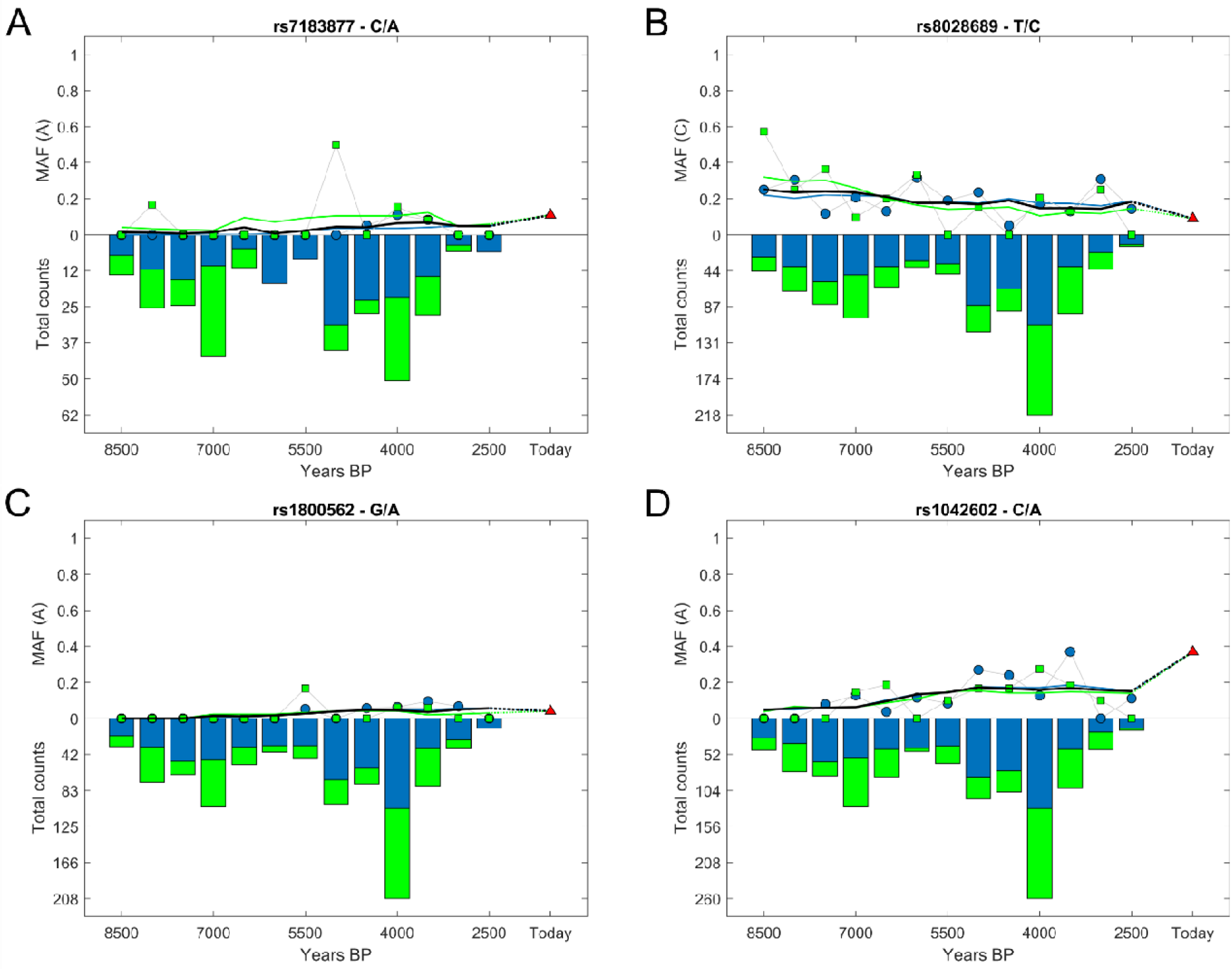
Temporal variation of the allele frequencies for four TIMs. Radiocarbon-dated samples (blue) and TPS dated samples (green) are shown. Bars show the number of individuals genotyped for each TIM. Lines show the changes in the minor allele frequency over time for each dataset. The black line shows the weighted average of the two MAF. A-B) TIMs rs7183877 (C/A) and rs8028689 (T/C) within the HERC gene are part of a haplotype spanning 166kB on chromosome 15, defined by 13 SNPs, found in 97% of all Caucasians with blue eyes, with values: rs7183877 (C) and rs8028689 (T). The other alleles (A for rs7183877 and C for rs8028689), seem to suggest brown eye colour ^82^. C) A TIM within the HFE gene associated with hemochromatosis. D) A TIM in the TYR gene. There is strong evidence that the rs1042602 A allele, associated with the absence of freckles, has been subject to positive selection in European populations ^83^.

## Tables legend

**Table S1**. (separate xlsx file) Ancient DNA dataset of 961 Eurasian individuals used to construct, calibrate, and test TPS. The database consists of radiocarbon-, archaeology-dated, and other/non-dated samples.

**Table S2**. (separate xlsx file) Temporal proportions obtained for 1) 300 ancient individuals for the determination of the 5 ancient temporal components; 2) 602 radiocarbon-dated individuals with 5 temporal components; 3) 300 ancient individuals for the determination of the 5 ancient and 3 modern temporal components.

**Table S3**. (separate xlsx file) 1) *p*-file output by ADMIXTURE with the allele frequencies associated with the 5 ancient components; 2) Corresponding Plink fam file; 3) Time series obtained for the TIMs.

**Table S4**. (separate xlsx file) TPS results for the ancient Eurasian database (Table S1) with eight temporal components. Filtering column P for “No” shows the 496 individuals in the reference panel, from which 400 random ones are selected in each run. These individuals were testing using the drop-one-individual approach and their results are shown in the “Drop-one-individual” sheet.

**Table S5**. (separate xlsx file) Ancient DNA dataset of 516 Central Asian individuals used to test TPS. The database consists of radiocarbon and archaeology dated samples.

**Table S6**. (separate xlsx file) TPS results for the 232 ancient Central Asian radiocarbon-dated individuals.

**Table S7**. (separate xlsx file) TPS results for 529 archaeology-dated Eurasian and Central Asian individuals.

**Table S8**. (separate xlsx file) Ancient DNA dataset of 143 Eurasian individuals used to test TPS. The database consists of radiocarbon-dated samples.

**Table S9**. (separate xlsx file) TPS results for 121 radiocarbon-dated Eurasian using various settings. TPS was applied to the samples using all the SNPs that overlapped with TPS SNPs or to their TIMs. Out of a total of ∼62k TIMs, on average each sample had ∼38k TIMs. TPS was then run on all these samples and two subsets: one where the test samples matched the training samples used to construct the TPS reference panel in country and period, and another one where the test samples did not match the training samples used to construct the TPS reference panel in country and period. Finally, the results are broken down by the number of SNPs available for analysis. TPS results include the number of samples analyzed, the mean and standard deviation of the accuracy, defined as the difference between TPS results and the mean radiocarbon date, the coefficient of determination (*r*^*2*^) and *t*-test’s *p*-value. All the results are plotted in Figs. S15-16.

**Table S10**. (separate xlsx file) Ancient DNA dataset of 110 Caucasus individuals used to test TPS. The database consists of mixed-dated samples.

**Table S11**. (separate xlsx file) TPS results for 110 mixed-dated Caucasus individuals.

**Table S12**. (separate xlsx file) A summary of TPS results obtained for 75 ancient Caucasus individuals using three reference panels, showing the median and 95% confidence intervals (for *N*>5).

**Table S13**. (separate xlsx file) TPS results for 67 Eurasian family relatives.

**Table S14**. (separate xlsx file) TPS results for 2117 modern-day individuals.

**Table S15**. (separate xlsx file) TPS results for four test-cases. TPS results were calculated using three reference panels.

## Data Legend

**Data S1. (separate zip file)**

Plink files with genetic data for the 961 ancient individuals and 147,229 SNPs used in this study.

**Data S2. (separate zip file)**

Plink files with the 8 temporal components generated in this study.

**Data S3. (separate zip file)**

Three excel file with the three reference panels for ancient populations and a fourth one for modern-day populations as used in this study.

## Acknowledgments

### Funding

This work was partially supported by the EPSRC Doctoral Training Partnership Grant EP/N509735/1 to U.E., MRC (MR/R025126/1) to E.E. and Crafoord foundation award to E.E.

### Author contributions

E.E. conceptualized the study. U.E., E.E. led the study. U.E., M.P. curated the data. U.E., G.H., G.A., A.M.D. analyzed the data. U.E., C.B., E.E. interpreted the results. U.E. wrote the TPS code. U.E., C.B., E.E. wrote the manuscript.

### Competing interests

E.E. Consults the DNA Diagnostics Center.

### Data and materials availability

All ancient data are available in the Supplementary Materials and in https://github.com/UEsposito/TPSpaper. TPS is available at https://github.com/UEsposito/TPStool.

## Supplementary Materials

Materials and Methods

Figures S1-18

Tables S1-16 (Supplementary Tables.zip)

Data S1 (Data S1.zip)

Data S2 (Data S2.zip)

Data S3 (Data S3.zip)

References (*1-64*)

## References

1 Taylor, R. E. & BarYosef, O. Radiocarbon Dating: An Archaeological Perspective, 2nd Edition. Radiocarbon Dating: An Archaeological Perspective, 2nd Edition, 1–404 (2014).

2 Higham, T. F. G., Jacobi, R. M. & Ramsey, C. B. AMS radiocarbon dating of ancient bone using ultrafiltration. Radiocarbon 48, 179–195 (2006).

3 MatisooSmith, E. & Horsburgh, K. A. DNA for Archaeologists. DNA for Archaeologists, 1–233 (2012).

4 Schaefer, N. K. & Shapiro, B. New middle chapter in the story of human evolution. Science 365, 981–982, doi:10.1126/science.aay3550 (2019).

5 Libby, W. F., Anderson, E. C. & Arnold, J. R. Age Determination by Radiocarbon Content: World-Wide Assay of Natural Radiocarbon. Science 109, 227–228, doi:10.1126/science.109.2827.227 (1949).

6 Be, M. M., Duchemin, B., Browne, E., Wu, S.C., Chechev, V., Helmer, R., & Schonfeld, E. Table of radionuclides comments on evaluations. (CEA Saclay, 1999).

7 Ramsey, C. B. Radiocarbon dating: Revolutions in understanding. Archaeometry 50, 249–275, doi:10.1111/j.1475-4754.2008.00394.x (2008).

8 Jacobi, R. M., Higham, T. F. G. & Ramsey, C. B. AMS radiocarbon dating of Middle and Upper Palaeolithic bone in the British Isles: improved reliability using ultrafiltration. J Quaternary Sci 21, 557–573, doi:10.1002/jqs.1037 (2006).

9 Brock, F., Higham, T., Ditchfield, P. & Ramsey, C. B. Current Pretreatment Methods for Ams Radiocarbon Dating at the Oxford Radiocarbon Accelerator Unit (Orau). Radiocarbon 52, 103–112, doi:Doi 10.1017/S0033822200045069 (2010).

10 Kromer, B. et al. Regional (CO2)-C-14 offsets in the troposphere: Magnitude, mechanisms, and consequences. Science 294, 2529–2532, doi:10.1126/science.1066114 (2001).

11 Alves, E. Q., Macario, K., Ascough, P. & BronkRamsey, C. The Worldwide Marine Radiocarbon Reservoir Effect: Definitions, Mechanisms, and Prospects. Rev Geophys 56, 278–305, doi:10.1002/2017rg000588 (2018).

12 Ascough, P., Cook, G. & Dugmore, A. Methodological approaches to determining the marine radiocarbon reservoir effect. Prog Phys Geog 29, 532–547, doi:10.1191/0309133305pp461ra (2005).

13 Biddle, M., Kjolbye-Biddle, B. in 13th Viking congress 45–96 (Oxbow, 1997 Nottingham, 2001).

14 Jarman, C. L., Biddle, M., Higham, T. & Ramsey, C. B. The Viking Great Army in England: new dates from the Repton charnel. Antiquity 92, 183–199, doi:10.15184/aqy.2017.196 (2018).

15 Dupree, L. Prehistoric research in Afghanistan (1959-1966). (American Philosophical Society, 1972).

16 Douka, K. et al. Direct radiocarbon dating and DNA analysis of the Darra-i-Kur (Afghanistan) human temporal bone. J Hum Evol 107, 86–93, doi:10.1016/j.jhevol.2017.03.003 (2017).

17 Allentoft, M. E. et al. Population genomics of Bronze Age Eurasia. Nature 522, 167–172, doi:10.1038/nature14507 (2015).

18 Olalde, I. et al. The Beaker phenomenon and the genomic transformation of northwest Europe. Nature 555, 190–196, doi:10.1038/nature25738 (2018).

19 Manning, S. W. et al. Fluctuating radiocarbon offsets observed in the southern Levant and implications for archaeological chronology debates. Proc. Natl. Acad. Sci. U. S. A. 115, 6141–6146, doi:10.1073/pnas.1719420115 (2018).

20 Korlevic, P., Talamo, S. & Meyer, M. A combined method for DNA analysis and radiocarbon dating from a single sample. Sci Rep-Uk 8, doi: ARTN 4127 10.1038/s41598-018-22472-w (2018).

21 Moorjani, P. et al. A genetic method for dating ancient genomes provides a direct estimate of human generation interval in the last 45,000 years. P Natl Acad Sci USA 113, 5652–5657, doi:10.1073/pnas.1514696113 (2016).

22 Meyer, M. et al. A high-coverage genome sequence from an archaic Denisovan individual. Science 338, 222–226, doi:10.1126/science.1224344 (2012).

23 Graur, D. et al. On the immortality of television sets: “function” in the Human genome according to the evolution-free gospel of ENCODE. Genome Biol. Evol. 5, 578–590, doi:10.1093/gbe/evt028 (2013).

24 Esposito, U., Das, R., Syed, S., Pirooznia, M. & Elhaik, E. Ancient Ancestry Informative Markers for Identifying Fine-Scale Ancient Population Structure in Eurasians. Genes (Basel) 9, doi:10.3390/genes9120625 (2018).

25 Alexander, D. H., Novembre, J. & Lange, K. Fast model-based estimation of ancestry in unrelated individuals. Genome Res 19, 1655–1664, doi:10.1101/gr.094052.109 (2009).

26 McLaren, W. et al. The Ensembl Variant Effect Predictor. Genome Biol. 17, 122, doi:10.1186/s13059-016-0974-4 (2016).

27 Myres, N. M. et al. A major Y-chromosome haplogroup R1b Holocene era founder effect in Central and Western Europe. Eur J Hum Genet 19, 95–101, doi:10.1038/ejhg.2010.146 (2011).

28 Underhill, P. A. et al. Separating the post-Glacial coancestry of European and Asian Y chromosomes within haplogroup R1a. Eur J Hum Genet 18, 479–484, doi:10.1038/ejhg.2009.194 (2010).

29 Underhill, P. A. et al. The phylogenetic and geographic structure of Y-chromosome haplogroup R1a. Eur J Hum Genet 23, 124–131, doi:10.1038/ejhg.2014.50 (2015).

30 Fu, Q. et al. The genetic history of Ice Age Europe. Nature 534, 200–205, doi:10.1038/nature17993 (2016).

31 Mathieson, I. et al. Genome-wide patterns of selection in 230 ancient Eurasians. Nature 528, 499–503, doi:10.1038/nature16152 (2015).

32 Salomonsen, B. Die Varby-funde. Ein Beitrag zur Kenntnis der altesten Trichterbecherkultur in Schonen. Acta Archaeologica, 55–95 (1970).

33 Skak-Nielsen, N. V. The neolithisation of Scandinavia: How did it happen? Adoranten (2004).

34 Paulsson, B. S. Scandinavian Models: Radiocarbon Dates and the Origin and Spreading of Passage Graves in Sweden and Denmark. Radiocarbon 52, 1002–1017, doi:Doi 10.1017/S0033822200046099 (2010).

35 Sørensen, T. in Materialities of Passing: Explorations in Transformation, Transition and Transience (ed Rasmussen AE Bjerrgaard P, Sørensen TF) 65–77 (Routledge, 2016).

36 Gasparri, S. Italia longobarda. Il regno, i Franchi, il papato. (Laterza, 2012).

37 Amorim, C. E. G. et al. Understanding 6th-century barbarian social organization and migration through paleogenomics. Nat Commun 9, 3547, doi:10.1038/s41467-018-06024-4 (2018).

38 Olalde, I. et al. Derived immune and ancestral pigmentation alleles in a 7,000-year-old Mesolithic European. Nature 507, 225–228, doi:10.1038/nature12960 (2014).

39 Wilde, S. et al. Direct evidence for positive selection of skin, hair, and eye pigmentation in Europeans during the last 5,000 y. Proc. Natl. Acad. Sci. USA 111, 4832–4837, doi:10.1073/pnas.1316513111 (2014).

40 Broushaki, F. et al. Early Neolithic genomes from the eastern Fertile Crescent. Science 353, 499–503, doi:10.1126/science.aaf7943 (2016).

41 Wang, W. Y. et al. Association Between Cartilage Intermediate Layer Protein and Degeneration of Intervertebral Disc A Meta-analysis. Spine 41, E1244–E1248, doi:10.1097/Brs.0000000000001749 (2016).

42 Günther, T. et al. Population genomics of Mesolithic Scandinavia: Investigating early postglacial migration routes and high-latitude adaptation. PLoS Biol. 16, e2003703, doi:10.1371/journal.pbio.2003703 (2018).

43 Mellars, P. A new radiocarbon revolution and the dispersal of modern humans in Eurasia. Nature 439, 931–935, doi:10.1038/nature04521 (2006).

44 Hofmanova, Z. et al. Early farmers from across Europe directly descended from Neolithic Aegeans. Proc Natl Acad Sci U S A 113, 6886–6891, doi:10.1073/pnas.1523951113 (2016).

45 Schiffels, S. et al. Iron Age and Anglo-Saxon genomes from East England reveal British migration history. Nat. Commun. 7, 10408, doi:10.1038/ncomms10408 (2016).

46 Lazaridis, I. et al. Genetic origins of the Minoans and Mycenaeans. Nature 548, 214–218, doi:10.1038/nature23310 (2017).

47 Amorim, C. E. G. et al. Understanding 6th-century barbarian social organization and migration through paleogenomics. Nat. Commun. 9, 3547, doi:10.1038/s41467-018-06024-4 (2018).

48 Damgaard, P. d. B. et al. 137 ancient human genomes from across the Eurasian steppes. Nature 557, 369–374, doi:10.1038/s41586-018-0094-2 (2018).

49 Fu, Q. et al. The genetic history of Ice Age Europe. Nature 534, 200–205, doi:10.1038/nature17993 (2016).

50 González-Fortes, G. et al. Paleogenomic Evidence for Multi-generational Mixing between Neolithic Farmers and Mesolithic Hunter-Gatherers in the Lower Danube Basin. Curr. Biol. 27, 1801-1810.e1810, doi:10.1016/j.cub.2017.05.023 (2017).

51 Haak, W. et al. Massive migration from the steppe was a source for Indo-European languages in Europe. Nature 522, 207–211, doi:10.1038/nature14317 (2015).

52 Harney, É. et al. Ancient DNA from Chalcolithic Israel reveals the role of population mixture in cultural transformation. Nat. Commun. 9, 3336, doi:10.1038/s41467-018-05649-9 (2018).

53 Jones, E. R. et al. The neolithic transition in the baltic was not driven by admixture with early European farmers. Curr. Biol. 27, 576–582 (2017).

54 Lipson, M. et al. Parallel palaeogenomic transects reveal complex genetic history of early European farmers. Nature 551, 368–372, doi:10.1038/nature24476 (2017).

55 Mathieson, I. et al. The genomic history of southeastern Europe. Nature 555, 197–203, doi:10.1038/nature25778 (2018).

56 Mittnik, A. et al. The genetic prehistory of the Baltic Sea region. Nat. Commun. 9, 442, doi:10.1038/s41467-018-02825-9 (2018).

57 Olalde, I. et al. The Beaker phenomenon and the genomic transformation of northwest Europe. Nature 555, 190–196, doi:10.1038/nature25738 (2018).

58 Unterländer, M. et al. Ancestry and demography and descendants of Iron Age nomads of the Eurasian Steppe. Nat. Commun. 8, 14615, doi:10.1038/ncomms14615 http://www.nature.com/articles/ncomms14615#supplementary-information (2017).

59 Esposito, U., Das, R., Syed, S., Pirooznia, M. & Elhaik, E. Ancient Ancestry Informative Markers for Identifying Fine-Scale Ancient Population Structure in Eurasians. Gene 9, 625 (2018).

60 Narasimhan, V. M. et al. The formation of human populations in South and Central Asia. Science 365, doi:10.1126/science.aat7487 (2019).

61 de Barros Damgaard, P. et al. The first horse herders and the impact of early Bronze Age steppe expansions into Asia. Science 360, doi:10.1126/science.aar7711 (2018).

62 Fernandes, D. et al. A genomic Neolithic time transect of hunter-farmer admixture in central Poland. Sci. Rep. 8, 1–11 (2018).

63 Fu, Q. et al. DNA analysis of an early modern human from Tianyuan Cave, China. Proc. Natl. Acad. Sci. USA 110, 2223–2227, doi:10.1073/pnas.1221359110 (2013).

64 Fu, Q. et al. Genome sequence of a 45,000-year-old modern human from western Siberia. Nature 514, 445–449, doi:10.1038/nature13810 http://www.nature.com/nature/journal/v514/n7523/abs/nature13810.html#supplementary-information (2014).

65 Fu, Q. et al. An early modern human from Romania with a recent Neanderthal ancestor. Nature, doi:10.1038/nature14558 (2015).

66 González-Fortes, G. et al. A western route of prehistoric human migration from Africa into the Iberian Peninsula. Proceedings of the Royal Society B: Biological Sciences 286, 20182288, doi:10.1098/rspb.2018.2288 (2019).

67 Jeong, C. et al. Bronze Age population dynamics and the rise of dairy pastoralism on the eastern Eurasian steppe. Proc. Natl. Acad. Sci. USA 115, E11248–E11255, doi:10.1073/pnas.1813608115 %J Proceedings of the National Academy of Sciences (2018).

68 Kilinç, G. M. et al. The demographic development of the first farmers in Anatolia. Curr. Biol. 26, 2659–2666 (2016).

69 Lazaridis, I. et al. Genomic insights into the origin of farming in the ancient Near East. Nature 536, 419–424, doi:10.1038/nature19310 (2016).

70 Lipson, M. et al. Ancient genomes document multiple waves of migration in Southeast Asian prehistory. Science 361, 92–95, doi:10.1126/science.aat3188 (2018).

71 Olalde, I. et al. The genomic history of the Iberian Peninsula over the past 8000 years. 363, 1230–1234, doi:10.1126/science.aav4040 %J Science (2019).

72 Raghavan, M. et al. Upper Palaeolithic Siberian genome reveals dual ancestry of Native Americans. Nature 505, 87–91, doi:10.1038/nature1273 http://www.nature.com/nature/journal/vaop/ncurrent/abs/nature12736.html#supplementary-information (2014).

73 Rodriguez-Varela, R. et al. Genomic Analyses of Pre-European Conquest Human Remains from the Canary Islands Reveal Close Affinity to Modern North Africans. Curr. Biol., doi:10.1016/j.cub.2017.09.059 (2017).

74 Schroeder, H. et al. Unraveling ancestry, kinship, and violence in a Late Neolithic mass grave. Proc. Natl. Acad. Sci. U. S. A., 201820210, doi:10.1073/pnas.1820210116 (2019).

75 Seguin-Orlando, A. et al. Genomic structure in Europeans dating back at least 36,200 years. Science 346, 1113–1118, doi:10.1126/science.aaa0114 (2014).

76 Sikora, M. et al. Ancient genomes show social and reproductive behavior of early Upper Paleolithic foragers. Science 358, 659–662, doi:10.1126/science.aao1807 (2017).

77 Ebenesersdottir, S. S. et al. Ancient genomes from Iceland reveal the making of a human population. Science 360, 1028–1032, doi:10.1126/science.aar2625 (2018).

78 Valdiosera, C. et al. Four millennia of Iberian biomolecular prehistory illustrate the impact of prehistoric migrations at the far end of Eurasia. Proc. Natl. Acad. Sci. USA 115, 3428–3433, doi:10.1073/pnas.1717762115 (2018).

79 Skourtanioti, E. et al. Genomic History of Neolithic to Bronze Age Anatolia, Northern Levant, and Southern Caucasus. Cell 181, 1158-1175.e1128, doi: https://doi.org/10.1016/j.cell.2020.04.044 (2020).

80 Genomes Project, C. et al. A global reference for human genetic variation. Nature 526, 68–74, doi:10.1038/nature15393 (2015).

81 Elhaik, E. et al. Geographic population structure analysis of worldwide human populations infers their biogeographical origins. Nat. Commun. 5, 1–12, doi:10.1038/ncomms4513 (2014).

82 Sulem, P. et al. Genetic determinants of hair, eye and skin pigmentation in Europeans. Nat Genet 39, 1443–1452, doi:10.1038/ng.2007.13 (2007).

83 Eiberg, H. et al. Blue eye color in humans may be caused by a perfectly associated founder mutation in a regulatory element located within the HERC2 gene inhibiting OCA2 expression. Hum Genet 123, 177–187, doi:10.1007/s00439-007-0460-x (2008).

